# Cell culture evolution of a HSV-1/VZV UL34/ORF24 chimeric virus reveals novel functions for HSV genes in capsid nuclear egress

**DOI:** 10.1101/2021.06.18.449081

**Authors:** Richard J. Roller, Tineke Hassman, Alison Haugo-Crooks

**Author notes:** Corresponding author, Department of Microbiology and Immunology, Carver College of Medicine, University of Iowa, 3-432 BSB, 51 Newton Road, Iowa City, IA 52242.

## Abstract

HSV and VZV are both members of the alphaherpesvirus subfamily, but are of different genera. Substitution of the HSV-1 UL34 coding sequence with that of its VZV homolog, ORF24, results in a virus that has defects in viral growth, spread, capsid egress, and nuclear lamina disruption very similar to those seen in a UL34-null virus despite normal interaction between ORF24 protein and HSV pUL31 and proper localization of the NEC at the nuclear envelope. Minimal selection for growth in cell culture resulted in viruses that grew and spread much more efficiently that the parental chimeric virus. These viruses varied in their ability to support nuclear lamina disruption, normal NEC localization and capsid de-envelopment. Single mutations that suppress the growth defect were mapped to the coding sequences of ORF24, ICP22 and ICP4 and one virus carried single mutations in each of the ICP22 and US3 coding sequences. The phenotypes of these viruses support a role for ICP22 in nuclear lamina disruption and a completely unexpected role for the major transcriptional regulator, ICP4, in capsid nuclear egress.

**Importance:** Interactions among virus proteins are critical for assembly and egress of virus particles, and such interactions are attractive targets for antiviral therapy. Identification of critical functional interactions can be slow and tedious. Capsid nuclear egress of herpesviruses is a critical event in the assembly and egress pathway and is mediated by two proteins that are conserved among herpesviruses, pUL31 and pUL34. Here we describe a cell culture evolution approach to identify other viral gene products that functionally interact with pUL34.

## Introduction

Herpesviruses form their genome-containing nucleocapsids in the cell nucleus, but accomplish final envelopment of those capsids at a cytoplasmic envelopment compartment composed of trans-Golgi and/or endosomal membranes. Movement of the capsid from the nucleoplasm to the cytoplasm is accomplished mainly by a sequential envelopment-de-envelopment process that is coordinated by a heterodimeric nuclear egress complex (NEC) composed of two conserved viral proteins (reviewed in (1)). In herpes simplex virus type 1 (HSV-1), the proteins are called pUL31 and pUL34. The nuclear egress process is an attractive therapeutic target, since the nuclear egress process is critical for assembly and egress of all herpesviruses, the proteins of the NEC are conserved among all herpesviruses, and the process is unlike anything yet observed in vertebrate cells.

The nuclear egress process can be broken down into multiple critical steps, most or all of which involve participation of pUL31 and pUL34, including disruption of the nuclear lamina, docking of genome-containing capsids, budding of capsid into the inner nuclear membrane (INM), scission of the inner nuclear membrane, fusion of the perinuclear virion envelope with the outer nuclear membrane (ONM), and release of the capsid into the cytoplasm. Completion of these steps in HSV-1infected cells requires recruitment of other viral and cellular factors to the nuclear envelope including protein kinases for disruption of the nuclear lamina and regulation of budding, the capsid itself, and ESCRT III components for scission of the perinuclear envelope virion (PEV) membrane (2-9).

The overall fold and arrangement of interaction surfaces of pUL31 and pUL34 homologs are highly conserved even between representatives of the alpha- and betaherpesviruses (10-14). Furthermore, the amino acid sequence determinants of the interactions that mediate heterodimerization are conserved within the sub-families. Schnee et al. reported interaction between pUL31 and pUL34 homologs from different members of the betaherpesvirus sub-family and between those of the alphaherpesviruses, pseudorabiesvirus (PrV) and HSV-1 (2).

In addition to their heterodimerization interaction, pUL31 and pUL34 heterodimers oligomerize during budding in vitro to form a hexagonal lattice (15, 16). This lattice is similar to that formed by the HSV NEC during crystallization, and it has been proposed that formation of this lattice drives curvature of the INM during budding (10, 14-16). Formation of hexamers, and interactions between adjacent hexamers in the lattice occur by interaction between the pUL34 and pUL31 proteins of adjacent heterodimers in the lattice (7, 10, 14).

Besides their interactions with each other, the NEC proteins have been reported to make interactions with many other viral and cellular proteins that contribute in some way to nuclear egress. The protein kinase pUS3 interacts with the NEC, phosphorylates both pUL31 and pUL34 and regulates both the nuclear lamina disruption and de-envelopment stages of nuclear egress (17-22). The NEC also recruits cellular kinases to the nuclear envelope partly by way of interaction with ICP34.5 and cellular protein p32 to contribute to lamina disruption (2-4). Additional interactions have been reported with viral proteins ICP22 and pUL47 (23, 24) and with cellular nuclear envelope proteins emerin and lamin A/C (25, 26). In most cases the mechanistic and quantitative significance of these interactions is unclear. Even in the case of the best studied functional interaction, that between the NEC and pUS3, the contribution of this functional interaction to the efficiency of nuclear egress is variable between cell types (27).

HSV-1 and varicella zoster virus (VZV) are both members of the alphaherpesvirus sub-family, but are members of different genera – the simplexviruses and the varicelloviruses, respectively (28). Although HSV and VZV share a host and overlapping cell tropisms within the host, their genera are thought to have diverged from each other at about the time of the mammalian radiation 65-70 million years ago (1). There is a striking 22% difference AT vs. GC base compositions of their genomes, with HSV being quite GC-rich and VZV being relatively GC-poor (29), and the median amino acid sequence identity between their shared proteins is about 35% (Table 1.). During the time since their divergence, it is likely that accumulating amino acid changes have rendered many important interactions between viral proteins incompatible between species, while interactions with host cell proteins might be conserved since the viruses share species and cellular tropisms. Based on these considerations we hypothesized that replacement of an HSV protein with its VZV homolog might result in a growth impaired virus that could then be used as a substrate for selection of mutations that restore critical interactions between viral proteins.

**Table 1.** Identity between genes common to HSV-1 and VZV^a^

| HSV-1 Gene | HSV protein length | VZV Gene | VZV protein length | Length of overlap | % identity in overlap |
| --- | --- | --- | --- | --- | --- |
| UL1 | 235 | ORF60 | 160 | 120 | 24.2% |
| UL2 | 244 | ORF59 | 305 | 240 | 53.3% |
| UL3 | 235 | ORF58 | 221 | 189 | 48.1% |
| UL4 | 199 | ORF56 | 244 | 169 | 36.7% |
| UL5 | 882 | ORF55 | 881 | 861 | 58.5% |
| UL6 | 640 | ORF54 | 769 | 612 | 45.1% |
| UL7 | 296 | ORF53 | 331 | 290 | 33.1% |
| UL8 | 750 | ORF52 | 771 | 678 | 32.0% |
| UL9 | 851 | ORF51 | 835 | 815 | 46.9% |
| UL10 | 341 | ORF50 | 435 | 231 | 39.4% |
| UL11 | 96 | ORF49 | 81 | 69 | 34.8% |
| UL12 | 626 | ORF48 | 551 | 489 | 37.2% |
| UL13 | 518 | ORF47 | 510 | 470 | 34.7% |
| UL14 | 219 | ORF46 | 199 | 191 | 34.6% |
| UL15 | 735 | ORF42 | 747 | 707 | 63.2% |
| UL16 | 373 | ORF44 | 363 | 347 | 30.0% |
| UL17 | 703 | ORF43 | 676 | 652 | 36.1% |
| UL18 | 318 | ORF41 | 316 | 311 | 43.7% |
| UL19 | 1374 | ORF40 | 1396 | 1358 | 53.5% |
| UL20 | 222 | ORF39 | 240 | 218 | 21.6% |
| UL21 | 535 | ORF38 | 541 | 467 | 35.8% |
| UL22 | 838 | ORF37 | 841 | 758 | 27.6% |
| UL23 | 376 | ORF36 | 341 | 327 | 31.2% |
| UL24 | 269 | ORF35 | 258 | 244 | 34.4% |
| UL25 | 580 | ORF34 | 579 | 559 | 46.0% |
| UL26 | 635 | ORF33 | 605 | 509 | 43.0% |
| UL27 | 904 | ORF31 | 931 | 862 | 48.5% |
| UL28 | 785 | ORF30 | 770 | 726 | 50.4% |
| UL29 | 1196 | ORF29 | 1204 | 1162 | 52.2% |
| UL30 | 1235 | ORF28 | 1194 | 1108 | 60.6% |
| UL31 | 306 | ORF27 | 333 | 300 | 53.7% |
| UL32 | 595 | ORF26 | 585 | 535 | 49.5% |
| UL33 | 130 | ORF25 | 156 | 120 | 40.1% |
| **UL34** | **275** | **ORF24** | **269** | **259** | **43.6%** |
| UL35 | 112 | ORF23 | 235 | 104 | 34.6% |
| UL36 | 3159 | ORF22 | 2763 | 2599 | 32.3% |
| UL37 | 1123 | ORF21 | 1038 | 957 | 30.6% |
| UL38 | 465 | ORF20 | 483 | 447 | 36.9% |
| UL39 | 1137 | ORF19 | 775 | 745 | 46.6% |
| UL40 | 340 | ORF18 | 306 | 306 | 60.8% |
| UL41 | 489 | ORF17 | 455 | 387 | 42.9% |
| UL42 | 488 | ORF16 | 408 | 376 | 25.5% |
| UL43 | 434 | ORF15 | 406 | 372 | 18.5% |
| UL44 | 511 | ORF14 | 560 | 461 | 29.3% |
| UL46 | 717 | ORF12 | 661 | 607 | 13.5% |
| UL47 | 693 | ORF11 | 793 | 603 | 25.5% |
| UL48 | 490 | ORF10 | 410 | 366 | 37.4% |
| UL49 | 301 | ORF9 | 302 | 239 | 31.0% |
| UL49A | 91 | ORF9A | 87 | 73 | 24.7% |
| UL50 | 371 | ORF8 | 396 | 335 | 28.12% |
| UL51 | 244 | ORF7 | 259 | 234 | 38.5% |
| UL52 | 1056 | ORF6 | 1083 | 958 | 53.3 |
| UL53 | 338 | ORF5 | 340 | 324 | 29.0% |
| UL54 | 512 | ORF4 | 452 | 445 | 31.7% |
| UL55 | 186 | ORF3 | 179 | 172 | 30.2% |
| UL56 | 234 | ORF0 | 129 | 123 | 33.3% |
| RL2 | 768 | ORF61 | 467 | 413 | 27.1% |
| RS1 | 1302 | ORF62 | 1310 | 806 | 57.8% |
| US1 | 420 | ORF63 | 278 | 264 | 30.7% |
| US10 | 303 | ORF64 | 180 | 146 | 34.2% |
| US9 | 90 | ORF65 | 102 | 86 | 26.7% |
| US3 | 481 | ORF66 | 393 | 365 | 44.4% |
| US7 | 390 | ORF67 | 354 | 312 | 29.8% |
| **Median identity** |  |  |  |  | **35.3%** |

<sup>a</sup>Protein sequences from HSV-1 strain (F) and VZV (SanDimas) were aligned using MUSCLE. Overlapping residues were scored for identity.

## Results

The VZV homolog of HSV UL34 is ORF24. The two proteins are of similar length, with 269 residues for VZV ORF24, and 275 for HSV-1 pUL34. Alignment of the ORF24 sequence with that of pUL34 using MUSCLE (30) indicates that the two proteins share ~44% amino acid identity overall, with most of the identical and highly similar residues located in the N-terminal 185 amino acids that roughly correspond to the globular domain (Table 1 and Figure 1A). There is less amino acid identity or similarity in the C-terminal region that is thought to be the disordered stalk with the transmembrane domain. The published crystal structure shows that 47 amino acids in pUL34 make contacts with pUL31 in the heterodimer (Figure 1A green (conserved) and red (non-conserved) highlighted residues). Interestingly, these amino acids are only 64% identical between the HSV and VZV pUL34 homologs. The published structure also shows that 13 residues in pUL34 make contact with pUL31 in the formation of NEC hexamers (Figure 1A aqua (conserved) and yellow (non-conserved) highlighted residues). These are much more highly conserved with 11 of 13 being identical and the two non-identical positions having biochemically similar side chains. Using the alignment shown in Figure 1A, the VZV ORF24 amino acid sequence can be threaded onto the HSV pUL34 crystal structure using SwissModel (3) with minimal deviations from the basic fold (Figure 1B), the most obvious of which is that, in the ORF24 model residues 32-35 are not predicted to form β strand β1. Overall the model suggests that the core fold of the two proteins is highly similar. Based solely on sequence conservation and structure modeling one might expect that replacement of UL34 by ORF24 would result in impaired replication due to significantly compromised heterodimer formation, but that normal budding might be facilitated by whatever heterodimers do form.

**Figure 1.**
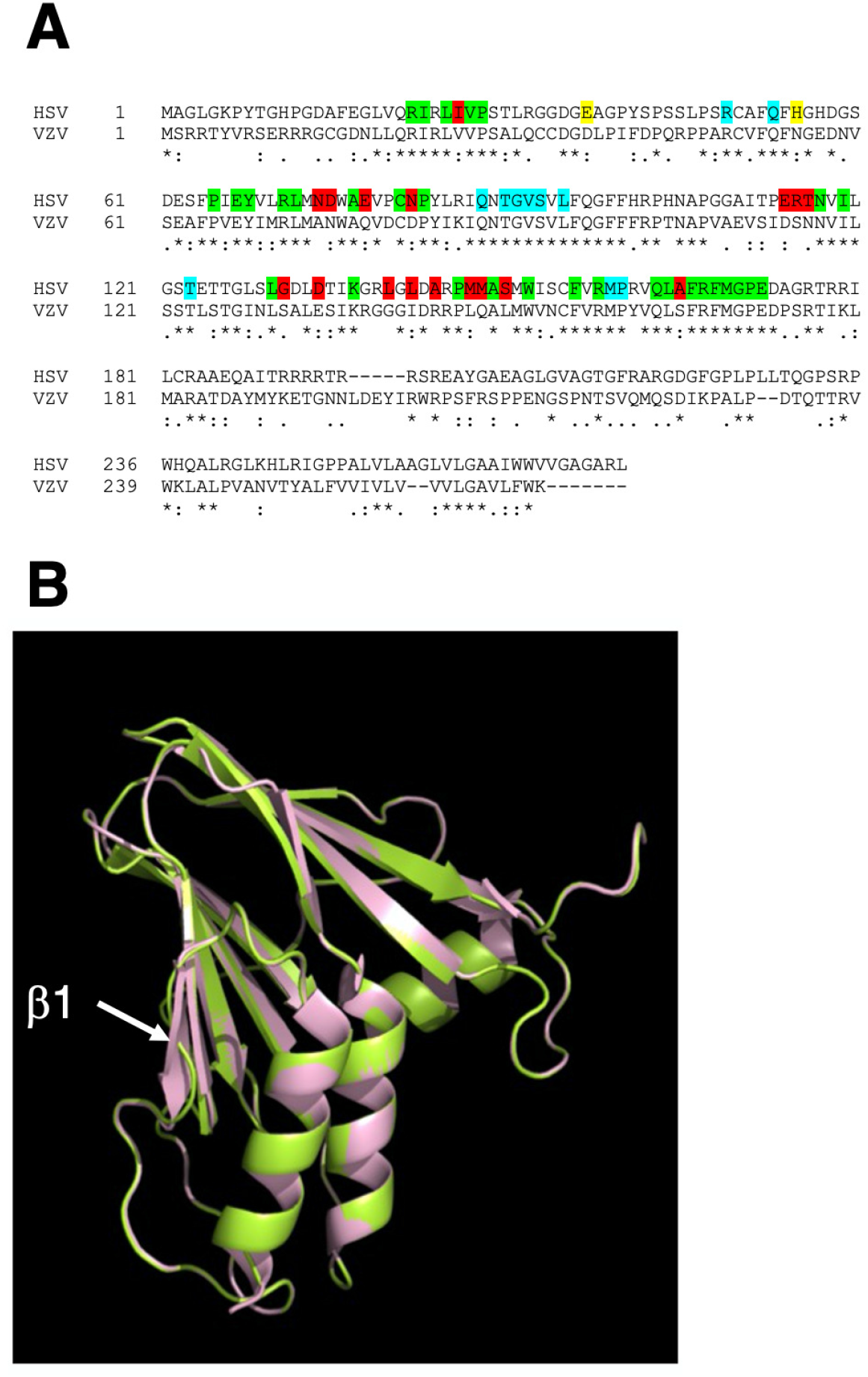
Sequence and structural alignment of HSV-1 pUL34 and VZV ORF24 proteins. (A) Sequence alignment of VZV ORF24 (top line) and HSV-1 pUL34 (bottom line) made using MUSCLE (30) and displayed in ClustalW format. Asterisks beneath the sequence lines indicate identity between sequences at that position; colons and periods indicate string and weak conservation of biochemical properties of side chains. Green and red highlighting in the HSV-1 sequence indicate conserved (green) and non-conserved (red) residues that participate in interactions with pUL31 in formation of the NEC heterodimer. Blue and yellow highlighting indicate conserved (blue) and non-conserved (yellow) residues that participate in interactions with pUL31 in adjacent NEC heterodimers in NEC hexamers. Identification of residues that participate in these interactions based on information provided in Bigalke and Heldwein (10). (B) Structural alignment of HSV-1 pUL34 (pink) and VZV ORF24 performed with SWISSMODEL (63) using the sequence alignment shown in (A).

To determine whether VZV ORF24 can functionally replace HSV-1 pUL34, a chimeric recombinant virus (called VZV ORF24) was constructed in which a full length, N-terminally FLAG epitope-tagged ORF24 coding sequence replaced most of the pUL34 coding sequence (Figure 2A, line 4). The sequences replaced were the same as those replaced by EGFP in the previously characterized UL34-null virus pRR1072(TK+) (31). Thus, VZV ORF24 would be expressed using the endogenous pUL34 promoter and regulatory elements. As a control, a recombinant virus encoding an N-terminally FLAG-tagged wild-type pUL34 (called FLAG-UL34) was also constructed (Figure 2A, line 3). Two independently derived isolates of the ORF24 replacement virus were constructed in order to ensure that any observed phenotype was due to the engineered genetic changes. Both viruses expressed proteins that reacted with anti-FLAG antibody at the size expected for pUL34 (Figure 2B). Surprisingly, the ORF24 protein runs slightly slower in SDS-PAGE than pUL34 even though it is five amino acids shorter. FLAG ORF24 is also expressed about 15-fold more poorly than FLAG-pUL34 in HSV-infected cells. Since ORF24 and pUL34 are expressed from the same promoter/regulatory elements in these viruses, this likely reflects differences in post-transcriptional regulation such as translation or stability.

**Figure 2.**
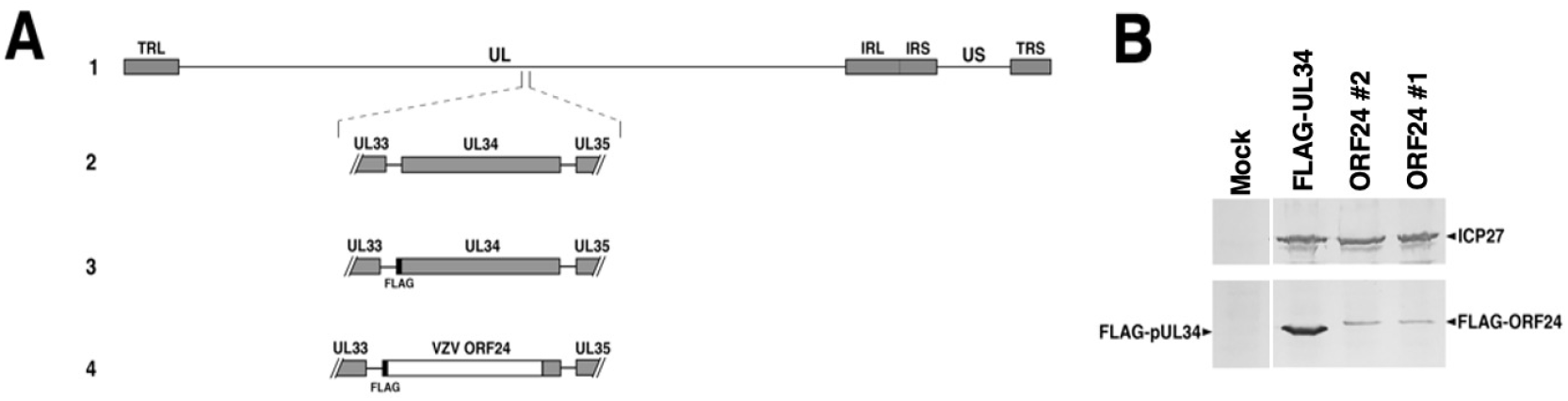
Construction of control and ORF24 replacement viruses and expression of ORF24 in infected cells. (A) (Line 1) Schematic diagram of the HSV-1 genome showing UL and US regions (thin lines) flaked by terminal and inverted repeats (gray boxes) (Line 2) expansion of the region of the UL34 gene showing the UL34 protein coding sequence (gray box). (Line 3) Structure of the UL34 region in the FLAG-UL34 control virus with the position of the FLAG tag (black box) indicated at the 5’ end of the UL34 coding sequence to express N-terminally-FLAG-tagged pUL34. (Line 4) Structure of the UL34 region in the ORF24 replacement virus. Most of the UL34 coding sequence from FLAG-UL34 is replaced by the complete ORF24 protein coding sequence (white bar). Some sequence from the 3’ end of the UL34 coding sequence was left to drive expression of UL35 (). (B) Expression of ORF24 from the replacement viruses. A digital image of an immunoblot of lysates from Vero cells infected with FLAG-UL34 and ORF24 virus isolates is shown. ICP27 (upper blot) is detected with anti-ICP27 antibody as an infection and loading control. FLAG-pUL34 and FLAG-ORF24 (lower blot) are detected with anti-FLAG antibody. All four lanes are from the same blot, but the mock lane was not adjacent to the other three.

To determine whether FLAG-ORF24 could support production of infectious virus, single-step growth of the wild-type HSV-1(F), UL34-null and ORF24 viruses was measured on Vero and UL34-expressing complementing cells (Figure 3A and B). On non-complementing Vero cells, the ORF24 replacement viruses grew with kinetics and final yield that were indistinguishable from a UL34-null virus, with peak virus production almost three log orders of magnitude lower than the wild type control. On HSV pUL34-expressing complementing cells, the UL34-null virus grew to wild-type levels, but the ORF24 replacement viruses produced significantly less (P<0.001 using ANOVA at 18 and 24 h.p.i.) virus than the WT and UL34-null controls, indicating that expression of ORF24 has a strong dominant negative effect on the function of wild-type HSV-1 pUL34.

**Figure 3.**
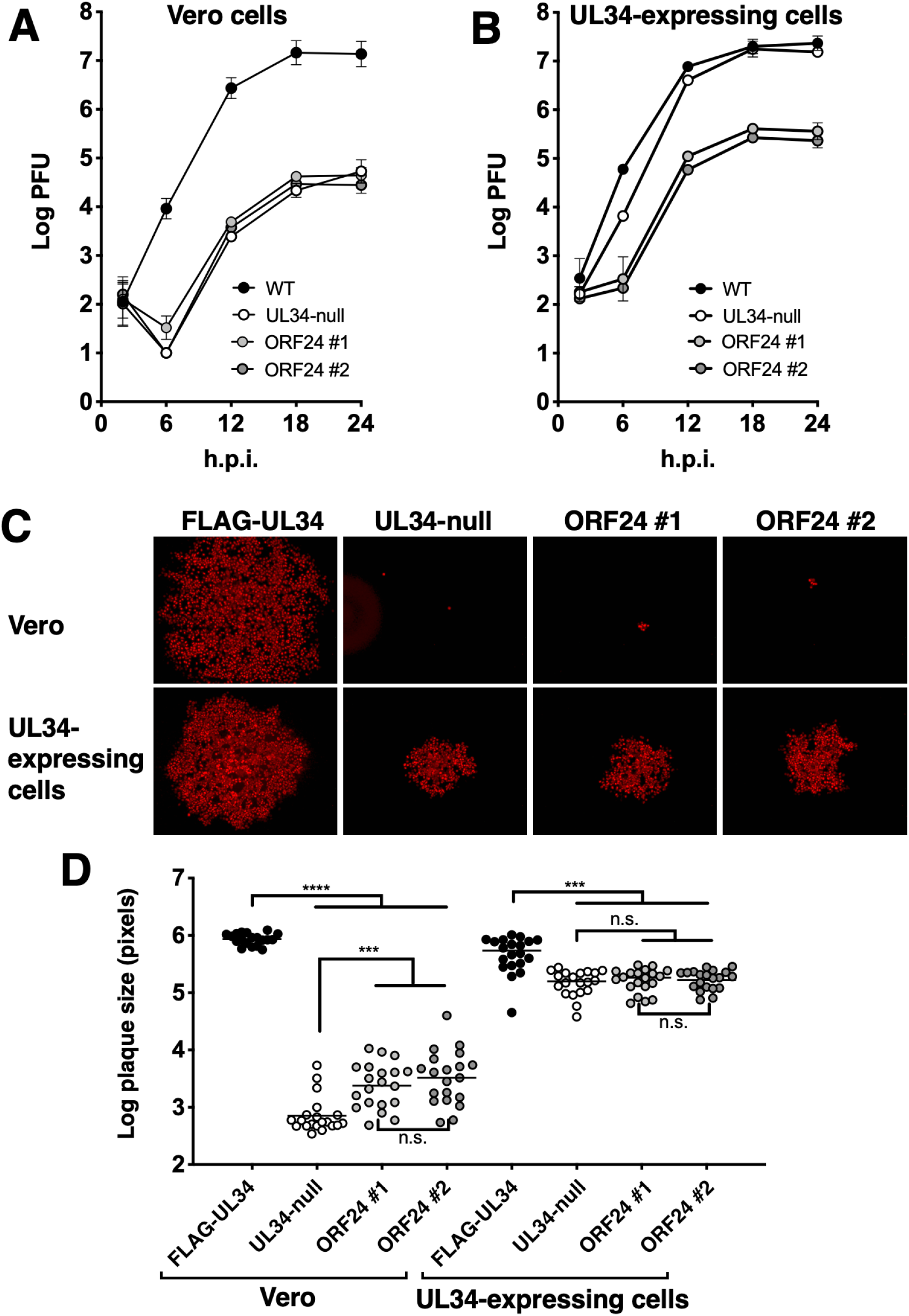
Growth and spread properties of ORF24 replacement viruses. (A) Single step growth of FLAG-UL34, UL34-null and ORF24 replacement viruses on Vero cells that express no pUL34 infected with 5PFU/cell of each virus. Each point represents the mean of three independent experiments and the error bars indicate the range of values. In some cases, the error bars are smaller than the point markers. (B) Same as (A), except cells that express wild-type pUL34 were infected. Statistical significance as indicated in the text was determined by performing a one-way ANOVA on log-converted values from single time points using the Method of Tukey for multiple comparisons implemented on GraphPad Prism. (C). Plaque formation in the presence of neutralizing antibody. Digital images of representative plaques formed by each of the indicated viruses on Vero cells (top row) or cells that express wild type pUL34 (bottom row). Plaques were visualized by immunofluorescent staining using antibody directed against the capsid scaffold protein. Plaques shown were of the median size from a sample of 20 randomly imaged plaques. (D) Plot of plaque areas measured from plaques visualized as in (C). Statistical significance was determined by performing a one-way ANOVA on log-converted values using the Method of Tukey for multiple comparisons implemented on GraphPad Prism. ***, P<0.001; ****, P<0.0001; n.s., not significant.

Measurements of plaques formed by the ORF24 replacement viruses on Vero cells in the presence of neutralizing antibody (Figure 3C and D) revealed that both of the replacement viruses formed plaques that were only slightly larger than those formed by UL34-null virus, and were much smaller than wild type. Interestingly, the plaques formed on UL34-complementing cells were the same size as those formed by UL34-null virus, indicating that there is no dominant negative effect on the spread function of HSV pUL34. The phenotypes of the two ORF24 isolates were indistinguishable in both single-step growth and spread assays on both cell types, indicating that the phenotypes were due to the ORF24 replacement.

pUL34 is thought to function by interaction with other viral and host cell protein partners. The failure of the ORF24 replacement virus to grow and the strong dominant negative effect of ORF24 expression suggested that ORF24 can interact with some subset of the partners of HSV pUL34 without forming a functional complex. The best characterized essential interaction partner of pUL34 is, of course, pUL31. In the absence of this interaction neither pUL31 nor pUL34 is properly recruited to the nuclear envelope in infected cells. To determine whether ORF24 is properly recruited to the nuclear rim during infection, Vero cells were infected with the FLAG-UL34 and ORF24 replacement viruses and then fixed and immunostained at 16 hours post infection (Figure 4A). The localization of ORF24 to the nuclear envelope in infected cells was indistinguishable from that of HSV pUL34 suggesting proper interaction with pUL31. To confirm that interaction can occur in the absence of other viral proteins, Vero cells were transfected with pUL31-FLAG and either HA-tagged pUL34 or HA-tagged ORF24 alone and in combination, and then fixed and immunostained at 48 hours post transfection. As expected, when transfected alone, pUL31 and HSV-1 pUL34 localized to the nucleoplasm, and to the nuclear envelope and to reticular cytoplasmic membranes, respectively (Figure 4B). When expressed alone, the localization of VZV ORF24 differed from HSV-1 pUL34, with no localization of ORF24 at the nuclear envelope, and formation of ORF24 puncta regularly distributed in the cytoplasm of the cell. However, when transfected together with UL31 (Figure 4C), both pUL34 and ORF24 co-localized with it at the nuclear rim and in internal nuclear structures, suggesting that ORF24 can interact with pUL31 in the absence of other viral factors. Interaction in transfected cells was confirmed by co-immunoprecipitation of HA-tagged pUL34 and HA-tagged ORF24 with FLAG-tagged pUL31 with similar efficiencies (Figure 4D)

**Figure 4.**
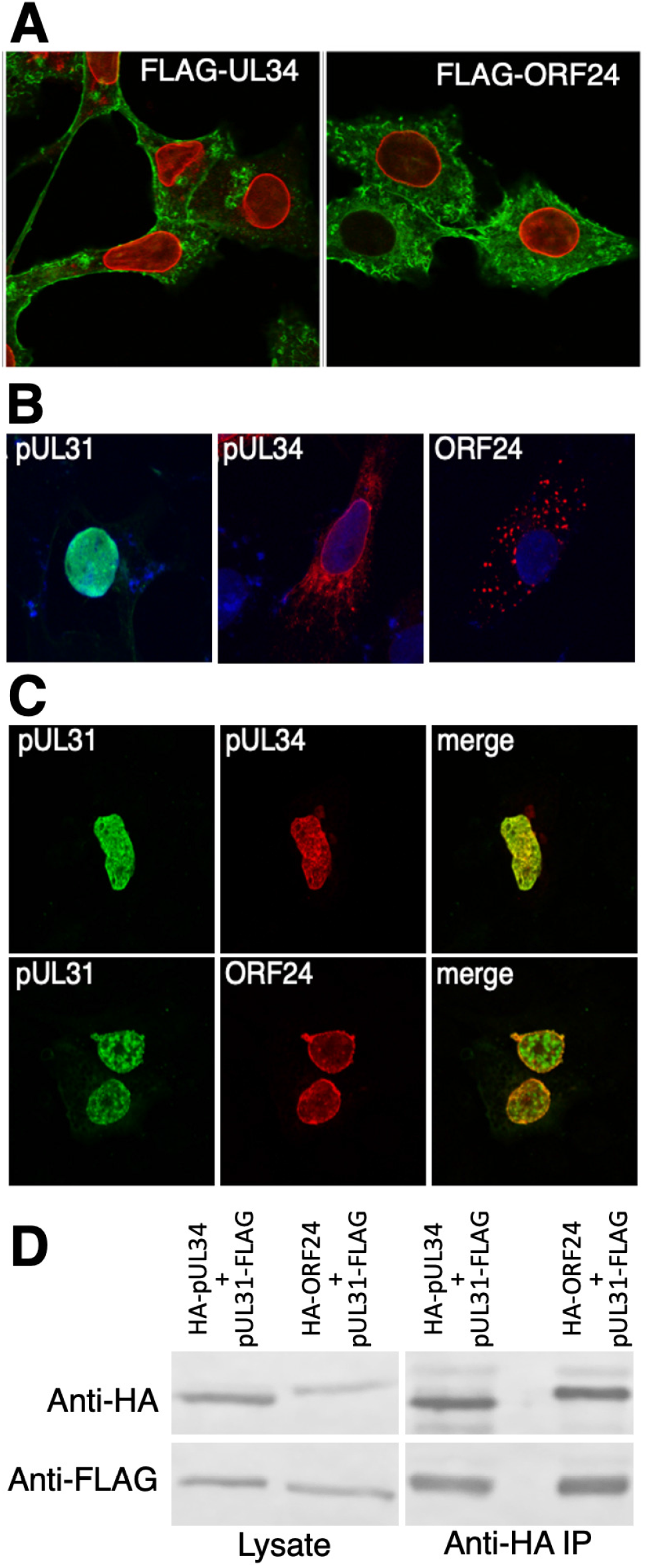
Interaction between VZV ORF24 and HSV-1 pUL31. (A) Localization of ORF24 and pUL34 in infected cells. Digital images of Z sections from Vero cells infected for 16 hours with the viruses indicated in each panel are shown. FLAG-pUL34 and FLAG-ORF24 were detected with anti-FLAG antibody and are shown in red. Filamentous actin is detected with fluorescently labeled phalloidin and is rendered in green. (B) Localization of pUL31-FLAG, HA-pUL34 and HA-ORF24 in individually transfected cells. Digital images of Z sections from Vero cells transfected with individual plasmids for 48 hours are shown. Nuclei are stained with Hoechst and rendered in blue. pUL31-FLAG was detected with anti-FLAG antibody and is rendered in green. HA-pUL34 and HA-ORF24 were detected with anti-HA antibody and are rendered in red. (C) Localization of FLAG-pUL31, HA-pUL34 and HA-ORF24 in co-transfected cells. Digital images of Z sections from Vero cells transfected with plasmids expressing pUL31-FLAG and either HA-pUL34 (top row) or HA-ORF24 (bottom row) for 48 hours are shown. pUL31-FLAG was detected with anti-FLAG antibody and is rendered in green. HA-pUL34 and HA-pUL31 were detected with anti-HA antibody and are rendered in red. (D). Co-immunoprecipitation of pUL31 FLAG with HA-pUL34 or HA-ORF24. HA-pUL34 and HA-ORF24 were purified from 293T cells transfected with the indicated plasmid combinations. Immunoblots of cell lysates (left) and magnetic-bead purified samples (right) were incubated with antibodies against the HA or FLAG epitope.

Since ORF24 interacts with HSV pUL31, one reasonable hypothesis to account for both its inability to function in virus replication and its dominant negative effect on HSV pUL34 function is that ORF24 fails to interact with other viral or cellular factors that are important for NEC function, and that its dominant negative effect results from interference with pUL34 function by sequestering pUL31 in defective NEC complexes. One approach to identification of other critical viral participants in NEC function is selection of extragenic suppressors of the growth phenotype. Six independent growth selections were performed by infection of non-complementing Vero cell cultures (Figure 5A). All six selections resulted in the development and spread of cytopathic effect. Single plaques were purified from each selection and then amplified on complementing cells to create stocks of each virus. Measurement of single-step growth at 16 hours post infection (Figure 5B) showed that all of the selected viruses (designated SUP1 through 6) significantly suppressed the growth defect of the parental VZV ORF24 virus, achieving >20-fold growth enhancement. Furthermore, all of the suppressor viruses restored growth to roughly the same degree. Notably, none of the SUP viruses restored growth to wild-type levels. All of the suppressors also significantly restored spread function, but without achieving wild-type plaque sizes (Figure 5C). The degree of spread enhancement among the suppressors was similar, except for SUP1, which formed significantly smaller plaques then the other suppressors.

**Figure 5.**
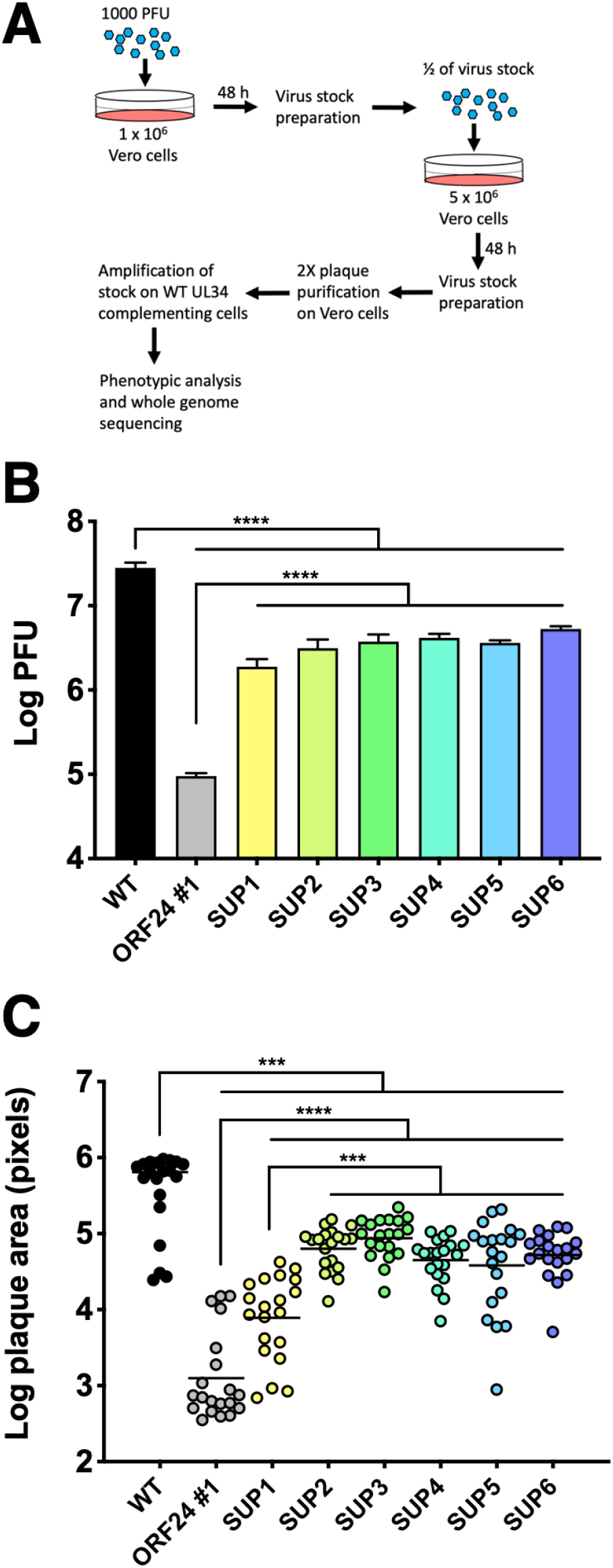
Growth and spread properties of suppressor viruses. (A) Virus plaque forming units produced 16 hours after infection of Vero cells with 5 PFU/cell of the indicated viruses. Bars represent the mean of three independent experiments. Each data point represents the mean of three independent experiments. Error bars in all panels represent the range of values obtained in three independent experiments. (B). Plaque sizes formed in the presence of neutralizing antibody. Plaque areas were measured 48 hours after infection at a low MOI. Each point represents a single plaque area. 20 plaques were measured for each condition. For both (A) and (B) statistical significance was determined by performing a one-way ANOVA on log-converted values using the Method of Tukey for multiple comparisons implemented on GraphPad Prism. ***, P<0.001; ****, P<0.0001.

The ORF24 replacement virus BAC was constructed on the pYE102bac background and we first sequenced both this BAC and the two ORF24 replacement virus BACs to create reference sequences for mapping Illumina reads from the suppressor viruses. A consensus sequence for pYE102bac was derived by mapping reads from the BAC onto a reference sequence created in silico using the published sequences of HSV-1 strain (F) and the BAC insertion described in Tanaka et al. (32, 33). Some sequence reads comprised of extended repeats could not be unambiguously aligned, including sequences at the genome termini, the sequence in RL that is proximal to the junction with UL, the junctions between the L and S components of the genome, and the sequence in US that is proximal to the junction with US. All other sequences, including all protein coding sequences could be unambiguously assigned. We observed 35 unique differences between pYE102bac and the published sequence of HSV-1 strain (F), including individual single-nucleotide polymorphisms (SNPs), a single-nucleotide insertion that produces an early frame-shift in the UL43 protein coding sequence, and a deletion between the UL29 and UL30 coding sequences that removes the long component origin of replication (Table 2). The sequences of the VZV ORF24 replacement BACs were identical to that of pYE102bac except for the expected replacement of UL34 sequences by ORF24. Illumina sequencing of the genomes from the parental VZV ORF24 #1 virus and the six suppressors gave average aligned read depths of ~400 at each position. Protein coding sequences all had aligned read depths >12 at each position. The sequence of the parental ORF24 #1 virus was identical to that of the BAC from which it was derived except for variations between reads in the number of residues in G/C runs in intergenic sequences. Many such intergenic sequences have G/C runs of 10-18 nucleotides and individual sequence reads from these areas showed variations of 1-3 nucleotides in the length of the run. These same differences were observed in the genomes of suppressor viruses. In addition, the suppressor virus genomes each contained interesting SNPs in protein coding sequences that were present in almost all reads from a given virus. Five of the six suppressor genomes (SUP1, 2, 4, 5, and 6) each had one SNP as compared to the parental genome and one of the six (SUP3) had two (Figure 6A). Each of these SNPs changed the encoded amino acid sequence. SUP4 and SUP6 had the same SNP in the ORF24 coding sequence itself that changed the glutamine at position 53 to arginine. The changes in SUPs 1, 2, 3, and 5 were all in other viral genes. SUP1 and SUP5 both had substitutions in the US1/US1.5 gene that encodes the immediate-early protein ICP22. SUP2 had a substitution in the coding sequence for the immediate early gene regulator ICP4. SUP3 had two substitutions, one in US1/1.5 and another in the coding sequence for the protein kinase pUS3.

**Table 2.** Sequence differences between HSV-1(F) and pYE102bac

| Nucleotide position in HSV-1(F)<sup>a</sup> | Gene or sequence region | Observed change<sup>b</sup> | Coding sequence change | Protein sequence change |
| --- | --- | --- | --- | --- |
| 2220 | ICP0 5' UTR | A to G | N/A |  |
| 9604 | UL1 | A to G | codon 111 GCA to GCG | silent |
| 15564 | UL6 | T to C | Codon 69 TTG to CTG | silent |
| 16429 | UL6 | C to T | Codon 457 GCT to GTA | A457V |
| 18753 | UL8 | G to A | Codon 552 CTG to TTG | silent |
| 19678 | UL8 | C to T | Codon 243 CCG to CCA | silent |
| 26904 | UL13 | C to T | Codon 507 TGT to TAT | C507Y |
| 34226 | UL15 exon 2 | T to C | Codon 568 TTG to CTG | silent |
| 35711 | UL18 | A to G | Codon 86 TGT to TGC | silent |
| 40404 | UL15 | C to T | Codon 15 GCG to ACG | A15T |
| 47555 | UL23 | G to C | Codon 55 GAC to GAG | D55E |
| 47695 | UL23 and UL24 | G to A | UL23 codon CAC to TAC | H9Y |
|  |  |  | UL24 codon GTG to GTA | silent |
| 47910 and 47911 | UL24 | CC to TG | Codon 86 GCC to GTG | A86V |
| 53638 | UL27 | G to T | Codon 691 CGC to CGA | silent |
| 61007 | UL29 | G to A | Codon 321 GAC to GAT | silent |
| 62302-62449 | OriL | deletion | N/A |  |
| 69681 | UL34 | G to A | Codon 45 AGC to AAC | S45N |
| 73069 | UL36 | T to C | Codon 2459 ATG to GTG | M2459V |
| 81439 | UL37 | G to A | Codon 850 TGC to TGT | silent |
| 82015 | UL37 | G to A | Codon 658 GAC to GAT | silent |
| 82806 | UL37 | G to A | Codon 395 CTG to TTG | silent |
| 82815 | UL37 | G to A | Codon 392 CTG to TTG | silent |
| 84529 | UL38 | T to C | Codon 35 TGC to CGC | C35R |
| 87759 | UL39 | C to T | Codon 474 GCT to GTT | A474V |
| 88762 | UL39 | C to T | Codon 808 TGC to TGT | silent |
| 94653-94654 | UL43 | C insertion | Codon 4 GTG to GCTG | Frame shift |
| 103968 | UL48 | G to A | Codon 335 TAC to TAT | silent |
| 104339 | UL48 | T to C | Codon 212 ACG to GCG | T212A |
| 107305 | UL50 | C to A | Codon 134 CCC to ACC | P134T |
| 115490 | UL55 | G to A | Codon 35 ATG to ATA | M35I |
| 116405 | UL56 | C to T | Codon 137 GCC to ACC | A137T |
| 123977 | ICP0 5' UTR | T to C | N/A |  |
| 127517 | ICP4 | G to A | Codon 1163 TGC to TGT | silent |
| 134689 | US2 | G to A | Codon 26 CTT to TTT | L26F |
| 137937 | US5/US6 intergenic | T to C | N/A |  |
| 139975 | US7 | A to G | Codon 110 GCA to GCG | silent |
| 150465 | ICP4 | C to T | Codon 1163 TGC to TGT | silent |

<sup>a</sup>Nucleotide positions are taken from (33).<sup>b</sup>Changes are noted on the top strand of the HSV-1(F) genome in prototype orientation.

**Figure 6.**
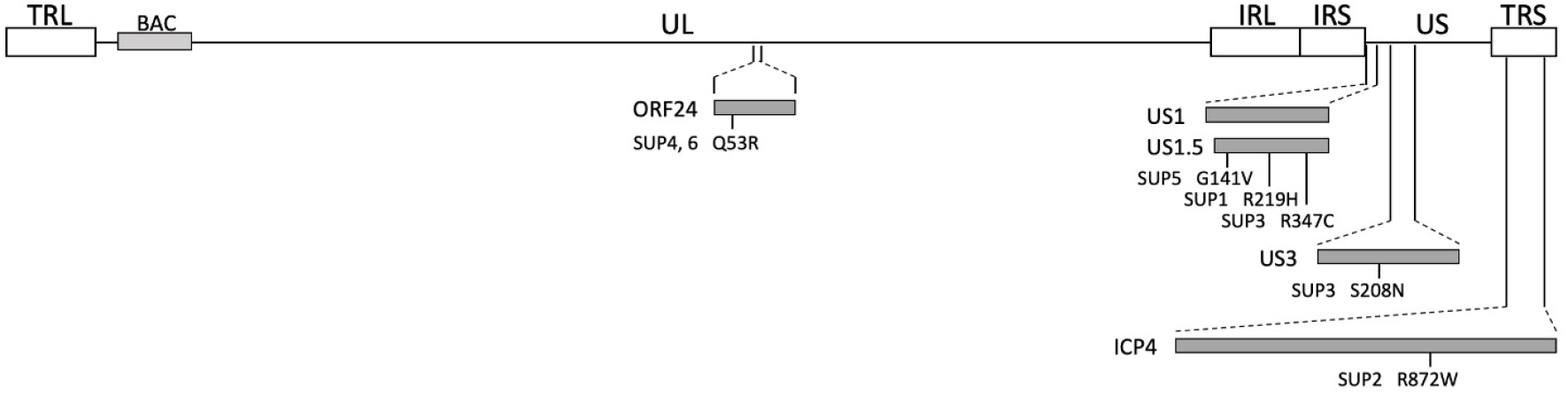
Location of mutations in ORF24 suppressor virus genomes. A schematic diagram of the HSV-1 ORF24 virus is shown showing the positions of terminal and inverted repeats (open boxes), the BAC vector sequence (gray box on the genome line), and protein coding sequences that have mutations in a SUP mutant (gray boxes projected below the genome line. The positions of amino acid changes are indicated by lines descending from the protein coding sequence box and the amino acid change is indicated below each line.

To determine what specific defects in nuclear egress might be suppressed by these mutations, protein expression, NEC localization, nuclear envelope disruption, and progress of nuclear egress by TEM were assessed. Expression of FLAG-ORF24 was similar in the parental and SUP viruses suggesting that the enhancement of growth seen in the SUP viruses was not due to greatly enhanced expression of ORF24 (Figure 7A). As shown above, localization of VZV ORF24 in cells infected by the parental virus is indistinguishable from that of pUL34 (Figure X). Interestingly, most of the suppressor mutants showed altered localization of the NEC in infected cells (Figure 7B). Only SUP5 showed roughly even staining of the nuclear envelope in most cells. SUP1 and 2, despite having mutations in different genes, showed the same localization pattern in which ORF24 staining was concentrated in blebs that extended from the nucleus into the cytoplasm. SUP3 showed a punctate distribution of ORF24 with protrusions into the nucleus reminiscent of that seen for US3-null or US3 catalytically dead mutants (). SUPs 4 and 6 both showed some distribution of ORF24 around the nuclear rim in all cells and in most cells (65% of 100 randomly selected cells observed for each mutant) showed a large concentration of staining off to one side of the nucleus.

**Figure 7.**
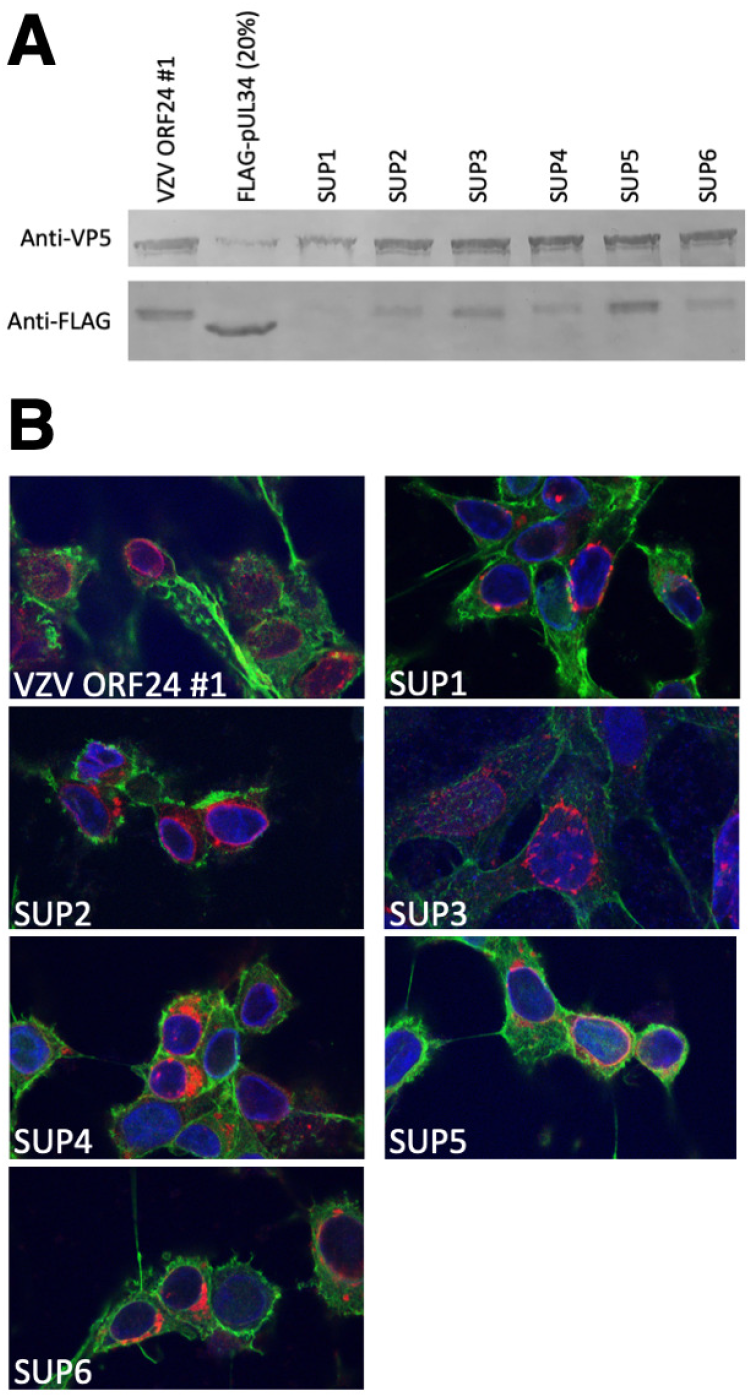
Expression and localization of suppressor mutant ORF 24. (A) Digital image of immunoblots of lysates from Vero cells infected with FLAG-pUL34 control virus, ORF24 #1 parental virus or ORF24 suppressor mutant viruses probed for VP5 as an infection and loading control or for FLAG to detect pUL34 or ORF24. The virus used for infection is indicated above the blot images. Equivalent volumes of whole cell lysate were loaded for each virus except for the FLAG-UL34 virus for which 20% of the volume was loaded to minimize oversaturation of signal. (B) Localization of ORF24 in SUP mutant virus infected cells. Digital images of Z sections from Vero cells infected for 16 hours with the viruses indicated in each panel are shown. FLAG-ORF24 was detected with anti-FLAG antibody and is shown in red. Nuclei were stained with Hoechst stain and are rendered in blue. Filamentous actin is detected with fluorescently labeled phalloidin and is rendered in green.

Previous studies using UL34, UL31, and US3 point mutations have shown that some mutations are associated with altered nuclear architecture and/or accumulation of specific nuclear egress intermediates, thereby providing information about the specific steps in nuclear egress that depend upon the correct amino acid sequence. In order to look for accumulation of such intermediates, Vero cells were infected with 5 PFU/cell for 20 h with wild-type, UL34-null, VZV ORF24 and SUP 1 through 5 viruses and then prepared for and examined by transmission electron microscopy (Figure 8). As expected, cells infected with control FLAG-UL34 virus showed multiple different assembly and egress intermediates, including intranuclear capsids, perinuclear enveloped virions (PEVs), cytoplasmic enveloped virions and mature virions on the cell surface (Figure 8A and Figure 9A). Cells infected with this virus also showed substantial deformation of the nuclear envelope indicating disruption of the nuclear lamina. In contrast, neither the UL34-null virus, nor the VZV ORF24 substitution viruses supported nuclear egress and very few extranuclear capsids were observed (Figure 8B and C and Figure 9A). All of the suppressor viruses showed some elevation in extranuclear capsids, but only SUP4 accumulated these to a level that was statistically significantly higher than the parental viruses (Figure 9A). The low levels of extranuclear capsids observed in the SUP virus-infected cells is consistent with the observation that these viruses replicate at least 10-fold less efficiently than the wild-type control (Figure 5). Interestingly, some (25%) of the sections of cells infected with the SUP4 virus showed accumulations of PEVs in extrusions of the outer nuclear membrane and ER that were concentrated to one side of the nucleus (Figure 8D). The similarity of this arrangement to that seen for pUL34 localization by immunofluorescence in SUP4 and SUP6 cells (Figure 7) suggests that these are the same structures and that the accumulation of pUL34 seen in immunofluorescence is due to the localized accumulation of perinuclear enveloped virions. Observation of these accumulations of perinuclear virions in only a subset of the cells examined in TEM likely results from sectioning of the infected cells in random planes, only some of which included the localized accumulations of PEVs.

**Figure 8.**
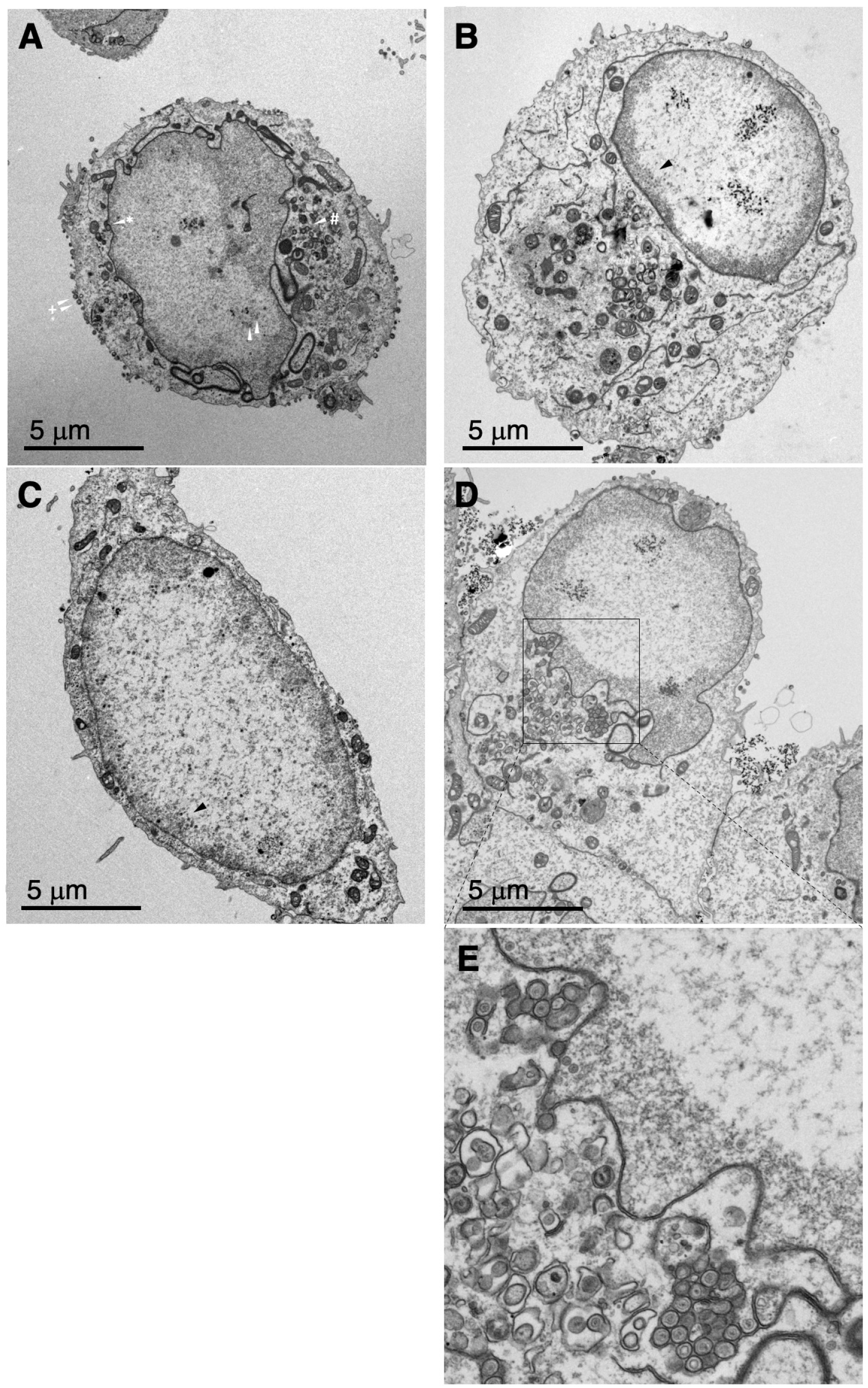
TEM analysis of recombinant and suppressor viruses. (A-D) 1000X magnification images of Vero cells infected for 20 hours with Flag-UL34 (A), UL34-null (B), VZV ORF24 #1 (C), and SUP4 (D) are shown. In (A), examples of different assembly intermediates are indicated with differently marked white arrowheads. Intranuclear capsids, white arrowhead; perinuclear enveloped virions, arrowhead with an asterisk; cytoplasmic enveloped virions, arrowhead with a pound sign; cell surface mature virions, arrowheads with plus sign. In (B) and (C), groupings of intranuclear capsids typical of nuclear egress-deficient virus infections are indicated with black arrowheads. The region in (D) containing groupings of perinuclear virions in extensions of the nuclear envelope/endoplasmic reticulum are boxed and expanded in (E).

**Figure 9.**
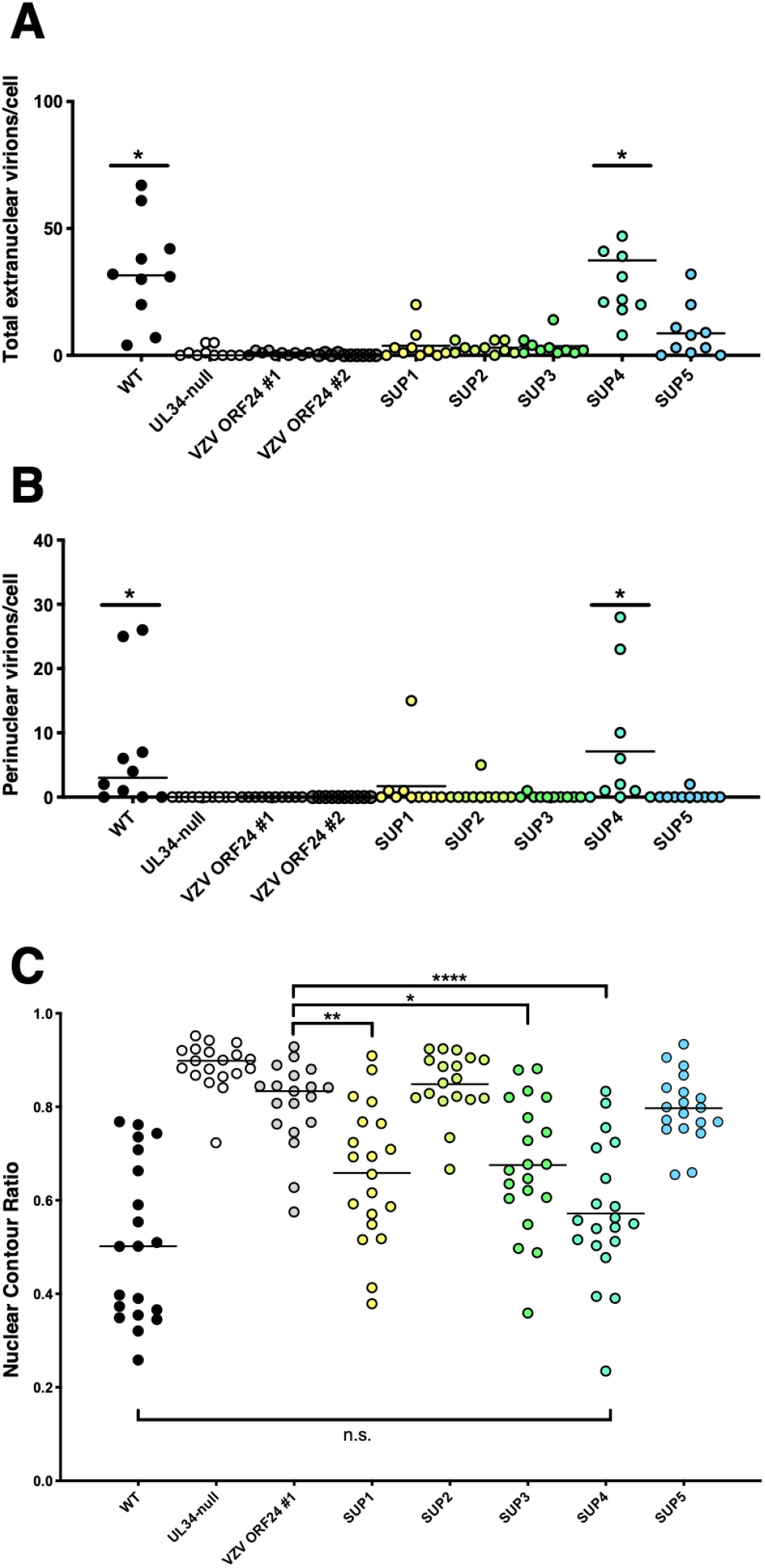
Quantitation of capsids in TEM images. (A) Nuclear envelopment measured as total capsids outside of the nucleoplasm (including perinuclear enveloped virions) counted in 1500X images of Vero cells infected for 20 h with the indicated viruses. Each point represents a section from a single cell. Ten sections were counted for each condition. (B) Counts of perinuclear enveloped virions per cell in the same sections analyzed in (A). (C) Quantitation of nuclear envelope deformation measured as nuclear contour ratio. Nuclear contour was defined in TEM sections of 20 cells at 1000X infected for 20 h with the indicated viruses. For (A) and (B) statistical significance was determined by performing a one-way ANOVA for comparison of all samples to VZV ORF24 #1 using the method of Dunnett implemented on GraphPad Prism. For (C) statistical significance was determined by performing a one-way ANOVA using the Method of Tukey for multiple comparisons of all samples to each other implemented on GraphPad Prism. *, P< 0.05; **, P<0.01, ***, P<0.001; ****, P<0.0001; n.s. not significant.

Disruption of nuclear architecture was measured using a nuclear contour ratio determined from TEM images as previously described (Figure 9B) (34). The nuclear contour ratio provides a measure of deviation of a contour from circularity. Higher ratios are indicative of contours close to smooth circularity, whereas lower ratios indicate increased convolution. As shown previously, the contour ratio of wild-type infected cells is much lower than that of uninfected cells and this change in contour is dependent upon UL34 function (34). The VZV ORF24 replacement virus is not significantly different from the UL34 null, indicating reduced ability to disrupt nuclear envelope architecture. Interestingly, the suppressors differ considerably in their ability to restore nuclear envelope disruption activity and this is not correlated with the site of the suppressor mutation. SUP1 (ICP22_R219H_), SUP3 (ICP22_R347C_/US3S_208N_) and SUP4 (ORF24_Q53R_) all restore nuclear envelope contour disruption to some degree, with SUP4 not significantly different from the wild-type control. SUP2 (ICP4_R872W_) and SUP5 (ICP22_G141V_) are not significantly better than the parental VZV ORF24 replacement virus.

To confirm that the SNPs observed in the SUP viruses were in fact the source of the suppressor phenotypes, individual recombinant mutant viruses containing these single mutations were constructed using the VZV ORF24 virus as parent. The genomes of each of the resulting viruses was sequenced to confirm the presence of the engineered mutations and the absence of other unintended mutations. Each of these viruses was tested for ability to suppress the single-step growth phenotype of the VZV ORF24 substitution (Figure 10).

**Figure 10.**
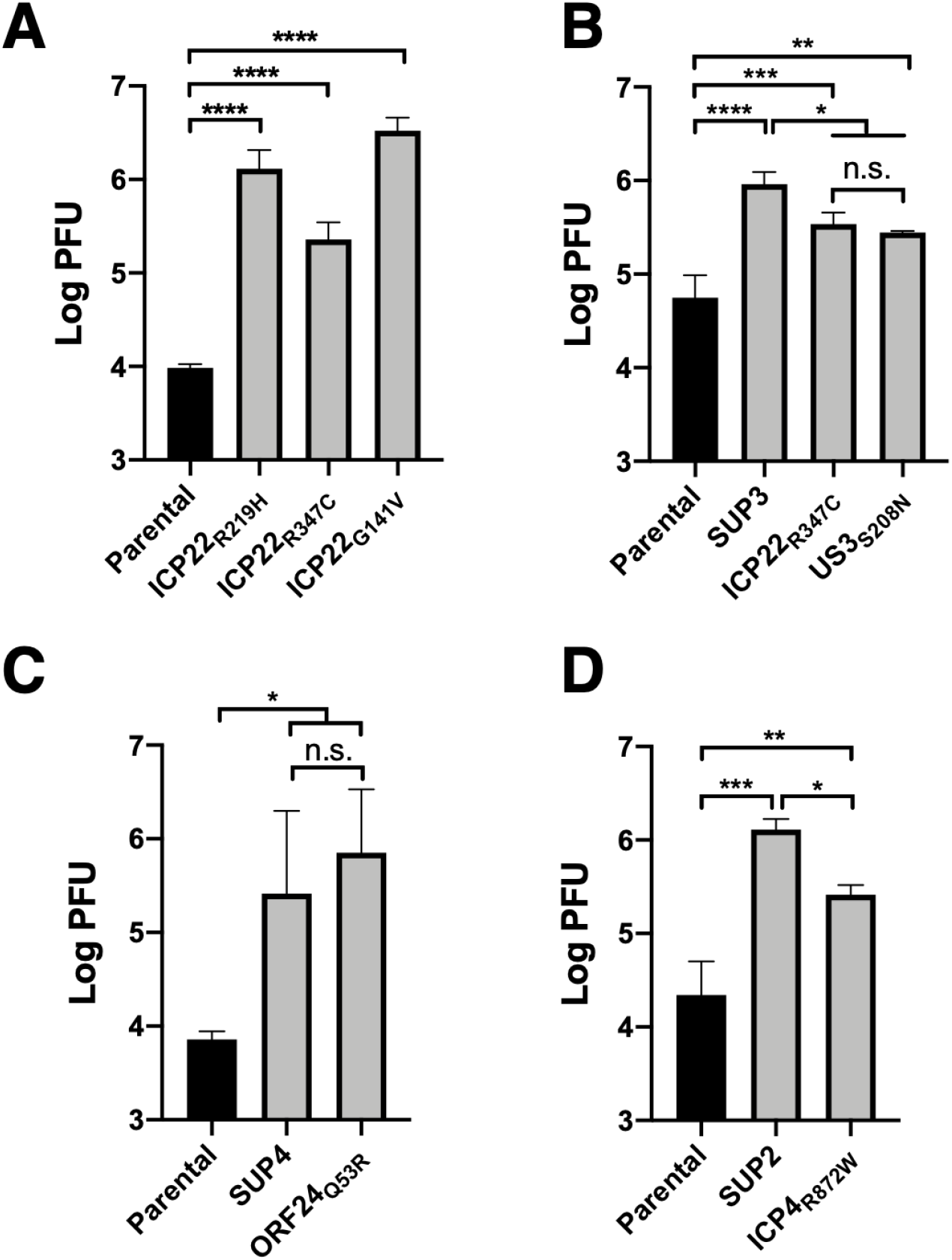
Recombinant suppressor single step growth. (A-D) Virus plaque forming units produced 16 hours after infection of Vero cells with 5 PFU/cell of the indicated viruses. (A) Growth of recombinant viruses with individual suppressor mutations in US1/ICP22. (B) Growth of recombinant viruses with individual mutations in ICP22 and US3 that make up the double mutant SUP3. (C) Growth of recombinant virus with the mutation in ORF24 contained in the SUP4 and SUP6 mutants. (D). Growth of recombinant virus with the ICP4 mutation contained in SUP2. Bars represent the mean of three independent experiments. Error bars in all panels represent the range of values obtained in three independent experiments. Statistical significance was determined by performing a one-way ANOVA on log-converted values using the Method of Tukey for multiple comparisons of all samples to each other implemented on GraphPad Prism. *, P< 0.05; **, P<0.01, ***, P<0.001; ****, P<0.0001; n.s. not significant.

Recombinant viruses carrying the mutations in the ICP22 coding sequence seen in SUP1 and SUP5 significantly suppressed the ORF24 growth defect and did so to a degree similar to that seen in the original suppressor viruses (Figure 10A). In contrast, the mutation in ICP22 seen in SUP3 improved growth compared to the ORF24 parent, but not to the same extent seen in the SUP3 suppressor virus, suggesting that this mutation alone was insufficient. To confirm this, single-step growth of recombinant viruses carrying each of the single SUP3 mutations were compared to the growth of the parental virus and to SUP3 itself (Figure 10B). Both of the individual mutations significantly improved growth compared to the parental ORF24 virus, but did so significantly more poorly than SUP3, suggesting that both mutations are necessary to achieve the full suppressive phenotype. The mutation in ORF24 found in the SUP4 and 6 suppressor viruses (Q53R) was sufficient to suppress the ORF24 growth defect to a degree seen similar to that seen in the SUP4 and 6 viruses (Figure 10C). The most surprising of the suppressor mutants was SUP2, which encodes a single amino acid substitution in ICP4. A recombinant virus carrying this single substitution on the ORF24 parent background was indeed able to significantly suppress the ORF24 growth defect (Figure 10D).

The localization pattern of the NEC in SUP3 (Figure 7) is reminiscent of that previously shown for recombinant viruses in which US3 is either deleted or catalytically inactivated, suggesting the possibility that the US3_S208N_ mutation results in loss of US3 expression or function. Expression of pUS3 was assessed by immunoblot of cells infected with the parental or SUP3 viruses and was observed to be the same (Figure 11A). To determine whether loss of pUS3 catalytic activity might suppress the ORF24 growth defect, two independent recombinant viruses were constructed using the VZV ORF24 BAC as parent in which the US3 gene was mutated to change lysine 220 to alanine (K220A). This substitution has been previously shown to ablate the kinase activity of pUS3 (). Introduction of the K220A mutation into US3 in recombinant mutants did not suppress the growth phenotype of the ORF24 virus (Figure 11B), suggesting that the S208N mutation in US3 is not simply a loss of function mutation. Indeed, loss of pUS3 kinase activity significantly depressed growth of the double mutant viruses and did so to the same degree in both viruses suggesting that the depressed growth was in fact due to the engineered mutation in pUS3. The presence of multiple mutations within the coding sequence for ICP22 was also consistent with the possibility that all of these were simple loss of function mutations. To test this hypothesis, recombinant viruses were constructed using the VZV ORF24 BAC as parent in which the US1 gene was completely deleted. In this case also, complete loss of ICP22 function did not suppress the ORF24 replacement growth phenotype and, in fact, resulted in poorer growth in the double mutant viruses (Figure 11C).

**Figure 11.**
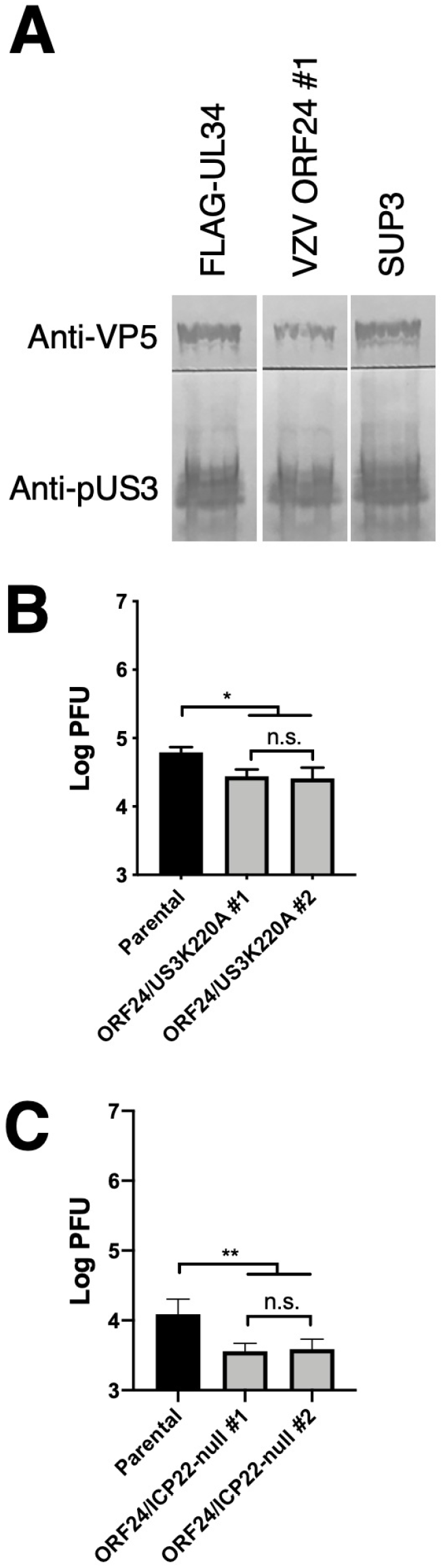
US3 expression and function in suppression of ORF24 growth defect. (A) A digital image of an immunoblot of lysates from Vero cells infected with FLAG-UL34 and ORF24 #1, and SUP3 viruses is shown. VP5 (upper blot) is detected with anti-VP5 antibody as an infection and loading control. pUS3 (lower blot) is detected with anti-pUS3 antibody. All three lanes are from the same blot, but were not adjacent in the original. The blot shown is representative of three independent experiments. (B) Growth of recombinant viruses with a catalytic inactivation mutation in pUS3 on the VZV ORF24 replacement virus background. (C) Growth of recombinant viruses in which the US1/1.5 gene has been deleted on the ORF24 replacement virus background. In (B) and (C), bars represent the mean of two independent experiments. Error bars in all panels represent the range of values obtained in three independent experiments. Statistical significance was determined by performing a one-way ANOVA on log-converted values using the Method of Tukey for multiple comparisons of all samples to each other implemented on GraphPad Prism. *, P< 0.05; **, P<0.01, ***, P<0.001; ****, P<0.0001; n.s. not significant.

## Discussion

The poor function of VZV ORF24 in HSV infection may result from two factors. The first is that the sequence divergence between HSV-1 and VZV has resulted in inability of ORF24 to make interactions that are critical for HSV pUL34 function. The importance of this factor is suggested by the ability to isolate suppressor mutants that recover some pUL34 functions without improving expression of the ORF24 protein. The other is that ORF24 is poorly expressed in the context of HSV1 infection, despite its expression being driven from the normal HSV UL34 5’ and 3’ regulatory sequences. This poor expression could derive from a number of factors, including poor translation of the VZV mRNA sequence due to the substantially different codon usage of HSV and VZV that derives from their very different GC contents, or low stability of the ORF24 protein. This low expression is not corrected in any of the suppressor mutants and this may explain why none of the suppressors achieves wild-type replication levels.

Functional suppression of the ORF24 defect might result from several effects of suppressor mutations. The most straightforward interpretation is that ORF24 cannot make specific interactions with other viral proteins that are necessary for capsid nuclear egress and that the suppressor mutations in those viral interaction partners restore one or more of those critical interactions. ICP22 and pUS3 have been previously shown to physically interact with the HSV-1 NEC and the presence of suppressor mutations in their coding sequences suggests the possibility that these mutations stabilize a physical interaction with the ORF24 NEC. In the same way, the presence of a suppressor mutation in ICP4 suggests that there may be a previously unsuspected physical interaction between ICP4 and the NEC. There are, however, other possible mechanisms of suppression including gain of function mutations that allow other viral factors to perform functions ordinarily performed by the wild-type NEC.

pUL31 and pUL34 make two types of critical interactions in accomplishing capsid nuclear egress. The first of these is the interaction that mediates formation of the pUL31/pUL34 heterodimer, and the second are the interactions between heterodimers that mediate hexameric array formation that is thought to drive curvature of the INM around the capsid. The heterodimerization interaction appears to occur efficiently between VZV ORF24 and HSV-1 pUL31. This is consistent with previous results of Schnee et al., showing interaction between the pUL31 and pUL34 proteins of members of the simplexvirus and varicellovirus genera (HSV-1 and PrV, respectively) (35). This is nonetheless surprising since amino acid side chains that make up the pUL31/pUL34 interaction surfaces in the heterodimer are not highly conserved. The second type of interaction can be subdivided into interactions between three different surfaces that meet in adjacent heterodimers within one NEC hexamer (intra-hexameric interactions) and between heterodimers that meet where hexameric rings interact with each other (inter-hexameric interactions (10, 15). Defects in these interactions manifest as budding defects in vitro and in infected cells (). The inter-hexameric interactions are dominated by interactions between adjacent pUL31 subunits of the NEC and might not be expected to be altered in the chimeric NEC. The intra-hexameric interactions, in contrast, are dominated by interactions between the pUL34 and pUL31 subunits of adjacent heterodimers and these interactions might be sensitive to the sequence differences between ORF34 and pUL34. We expected, therefore, to isolate suppressor mutations in both ORF24 and pUL31 that might compensate for destabilization of this interaction. Specifically, we have previously isolated a single R229L substitution mutant in pUL31 that suppresses multiple mutations that cause defects in intra-hexameric interaction (34, 36). We were surprised to find no suppressor mutations in the UL31 gene in this screen. The occurrence of suppressor mutations within the VZV ORF24 coding sequence itself as seen in SUPs 4 and 6 was unsurprising, but mutation at Q53 was unexpected because this residue does not vary between HSV pUL34 and VZV ORF24 and, in fact, is conserved among all of the mammalian alphaherpesviruses. In the HSV NEC hexamers observed in crystals (10), the side chain of Q53 is at the interface between the pUL34 of one heterodimer and the pUL31 of its neighboring heterodimer where it can participate in the intra-hexameric interaction by making hydrogen bond interactions with the main chain carbonyl oxygen and side-chain sulfur of M112 of pUL31 (Figure 12A and B). The substitution with arginine seen in SUPs 4 and 6 introduces a longer side chain, and a SWISS-MODEL prediction of the position of that side chain suggests that it might make more extensive interactions with the side chain of pUL31 M112 (Figure 12C and D) thereby stabilizing hexamer formation and promoting NEC budding function.

**Figure 12.**
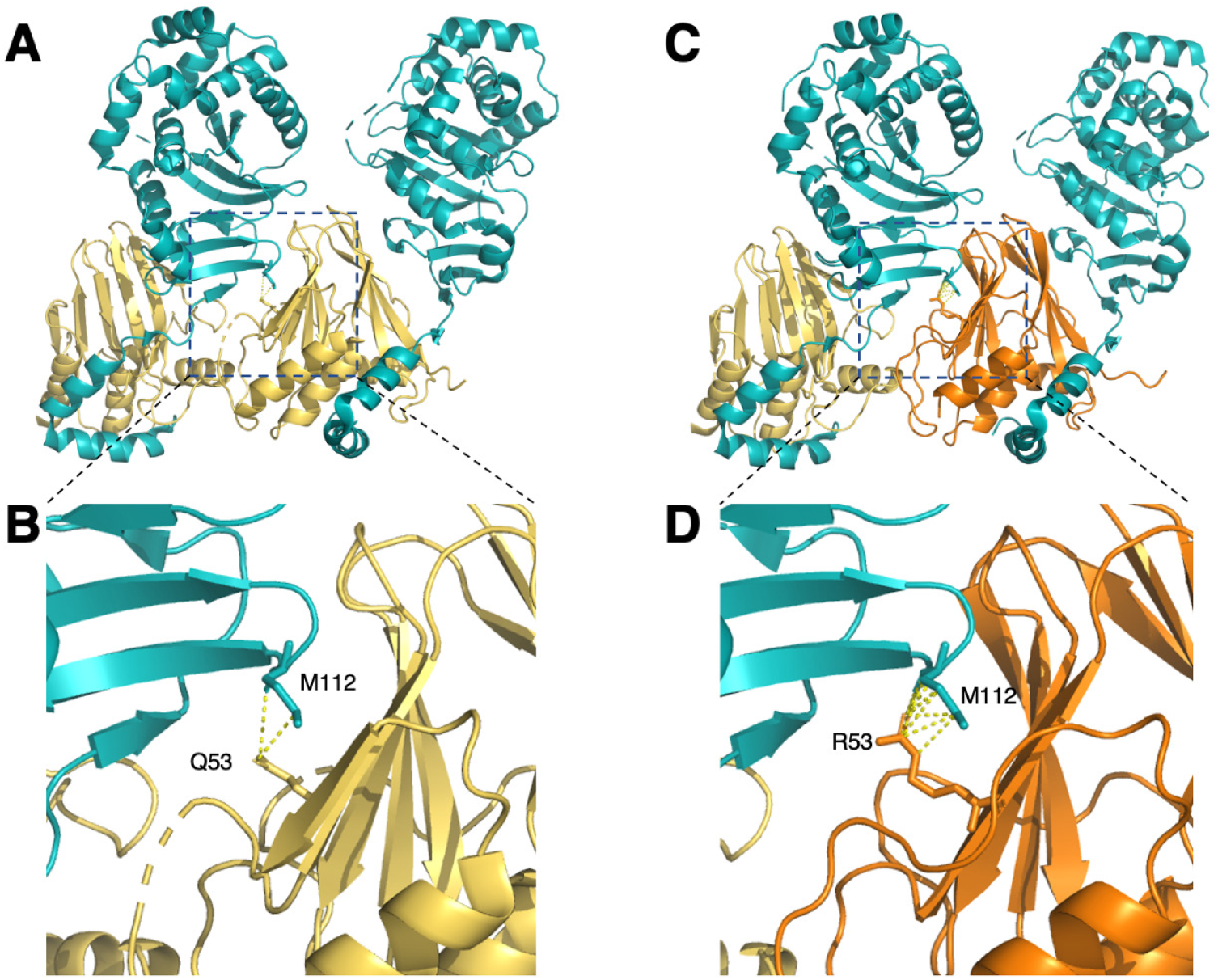
Alteration of interaction between ORF24 and pUL31 in the ORF24 Q53R mutant. (A) Ribbon diagram of the structure of two adjacent NEC heterodimers in a hexamer derived from entry 4ZXS in the Research Collaboratory for Structural Bioinformatics Proteins Data Bank (RCSB PDB) rendered using MacPyMol. pUL34 molecules are colored gold and pUL31 molecules are colored teal. Residues Q53 of pUL34 and M112 from pUL31 from adjacent heterodimers are depicted as sticks with hydrogen bonds shown as dotted yellow lines. (B), Expansion of boxed region in (A). (C) Ribbon diagram of two adjacent NEC heterodimers in which the pUL34 of the right heterodimer is replaced by a model of ORF24Q53R derived using homology modeling with SWISSMODEL using the same alignment as in Figure 1. pUL31 is shown in teal, pUL34 in yellow, and the modeled ORF24 in orange. Residues R53 of ORF24Q53R and M112 from pUL31 from adjacent heterodimers are depicted as sticks with hydrogen bonds and dipole interactions shown as dotted yellow lines.

ICP22 is an immediate early gene product and is conserved among the alphaherpesviruses, but is not found in the beta- and gammaherpesviruses (37). Its first identified function was in regulation of viral gene expression, and ICP22 has been reported to both facilitate expression of a subset of the viral late genes and to inhibit transcriptional elongation on some host and virus genes (38-42), but additional functions have been found (38-42). The isolation of three different suppressor viruses that carry mutations in ICP22 strongly supports an important role for ICP22 in HSV nuclear egress, as originally suggested by Muruzuru et al. (24). While that study did not address the mechanism of ICP22 action in nuclear egress, experiments here suggest a role for ICP22 in multiple processes including the disruption of nuclear architecture that accompanies capsid egress. Replacement of UL34 with ORF24 results in failure to disrupt the nuclear lamina as indicated by changes in nuclear contour during infection. Several of the suppressor viruses, including two of the ICP22 mutants (SUP1 and SUP3), substantially restore lamina disruption, suggesting that ICP22 helps to mediate this process. It seems likely that ICP22 has at least one additional function in nuclear egress, since SUP5 grows as well as SUPs 1 and 3 despite no significant improvement in disruption of nuclear architecture. One of the functions of ICP22 is to facilitate the formation of nuclear foci called virus-induced, chaperone-enriched (VICE) domains and ICP22 itself has recently been shown to have homology to a family of cellular chaperones (43, 44). One hypothesis to explain its participation in phenotypic suppression is that mutations in ICP22 enable it or other VICE domain chaperones to more effectively chaperone the NEC into a fully functional conformation. It has previously been observed that mutations that severely disrupt interaction between pUL31 and pUL34 when those proteins are expressed ectopically have a lesser effect in infected cells suggesting that some viral factor can facilitate their interaction (25). It is tempting to speculate that this factor might be ICP22 acting in its capacity as a chaperone.

The mechanism of phenotypic suppression by the S208N mutation in pUS3 is unclear. S208 is at the border between the N-terminal regulatory domain and the catalytic domain of the kinase. Mutation at this position might, therefore, have either appositive or negative effect on pUS3 kinase activity. The punctate distribution of the ORF24 in SUP3-infected cells is consistent with impairment or loss of kinase activity, but complete ablation of pUS3 kinase activity by mutation of the kinase domain invariant lysine at position 220 does not suppress the ORF24 growth defect. This suggests that possibility that the S208N substitution might partially inhibit kinase activity. Loss of pUS3 activity promotes NEC aggregation and this may reflect an ability of pUS3 to regulate the self-association of NEC heterodimers into hexameric arrays that promotes membrane curvature in envelopment (27). It is possible that the ORF24/pUL31 chimeric NEC is deficient in this self-association activity and that a mutation that inhibits pUS3 kinase activity might favor array formation and partially restore envelopment. It is also notable that the S208N substitution changes a potentially phosphorylatable residue to one that is non-phosphorylatable and many protein kinases are activated by phosphorylation in their regulatory domains. S208 is not, however, part of any previously characterized motif for phosphorylation by viral or host kinases.

The herpesvirus NEC coordinates or directly carries out multiple tasks in the nuclear egress process, including disruption of the nuclear lamina, capsid docking, membrane curvature, membrane scission, de-envelopment fusion and capsid release to the cytoplasm. The phenotypes of the various suppressors suggests that the ORF24/UL31 chimeric NEC is at least partially deficient in its ability to carry most, or all, of these tasks. The chimeric NEC is unable to mediate normal disruption of nuclear architecture as indicated by the smooth contour of nuclei in cells infected with the parental chimeric virus. However, suppressor mutants that substantially restore disruption of nuclear architecture (SUP1, 3, and 4) do not restore wild-type levels of replication, suggesting that the chimeric NEC is deficient in its ability to support later steps as well. Furthermore, SUPs 1, 3, and 4 differ in their nuclear egress deficiencies. Specifically, SUP1 and SUP3 do not accumulate specific intermediates in the nuclear egress process, whereas SUP4 accumulates primary enveloped virions in evaginations of the outer nuclear membrane in the cytoplasm. This suggests that SUPs 1 and 3 are deficient enough in their ability to mediate envelopment of capsids at the inner nuclear membrane that perinuclear virions do not accumulate. SUP4, on the other hand, may envelop capsids relatively efficiently, but are deficient in either de-envelopment membrane fusion or capsid release into the cytoplasm.

The most surprising finding from suppressor sequencing was the occurrence of a mutation that suppresses the ORF24 growth defect within the gene encoding the major transcription regulator, ICP4. ICP4 is an essential immediate early gene product that activates transcription from early and late gene promoters and can repress transcription from its own and other IE gene promoters (45). It has not been previously reported to have any role in capsid assembly or egress, although it is interesting to note that pUS3 co-purifies with ICP4 in tandem-affinity purification (46), suggesting an interaction with at least one of the viral factors that mediates nuclear egress. ICP4 is an elongated homodimeric protein whose structure is only partially characterized (47, 48). The structure of its DNA binding domain has been determined (48), and regions required for nuclear localization, homodimerization, induction of early gene expression and DNA replication and subsequent late gene expression have been roughly defined (45, 49-54). ICP4 R872 is within the C-terminal 40% of the protein that is required for efficient DNA replication and late gene expression, but not for expression of early genes (49, 50), and is found in a relatively highly conserved region of the protein that stretches from residue 798-876. R872 itself is completely conserved among the mammalian alphaherpesviruses including VZV, suggesting importance for some critical function of ICP4. The lack of any previous evidence for participation of ICP4 in nuclear egress might be attributable to its essential role for viral gene expression. While it is possible that the effect of ICP4 mutation on viral growth in the VZV ORF24 virus is an indirect effect of changes in gene expression, this mutation does not enhance the expression of ORF24 itself (Figure 7).

Another surprising result of these studies is that mutations that suppress the ORF24 defects do not create amino acid substitutions that make the HSV protein more “VZV-like.” Indeed, three of the suppressor substitutions (SUP1, SUP2, and SUP4/6) alter amino acids that are conserved between VZV and HSV-1 in ICP22, ICP24, and ORF24, respectively. The remaining substitutions affect non-conserved residues, but do not change them to correspond to their VZV counterpart residues. The substitutions in highly conserved residues might be expected to impair normal function of the mutant protein, and whether this can be observed on an otherwise wild-type HSV-1 background will require further study.

## Materials and Methods

### Cells and Viruses

All viruses used in these studies were derived from the pYEbac102 clone of the HSV-1 strain (F) genome in the bacterial strain GS1783 (kind gift of G. Smith) (32, 55). Viruses were propagated on Vero tUL34 CX cells that express HSV-1 pUL34 under the control of its own promoter regulatory sequences (25). Vero tUL34 CX cells were propagated in DMEM high glucose supplemented with 5% fetal bovine serum and the antibiotic penicillin and streptomycin.

### Recombinant virus constructions

Recombinant viruses were constructed by BAC recombineering as described by Tischer et al. (56, 57). The primers used for recombinant virus construction are shown in Table 3. For construction of a chimeric recombinant in which FLAG-tagged VZV ORF24 is expressed in place of pUL34 (hereafter referred to as FLAG-VZV-24), a cassette containing FLAG-ORF24 and a gentamycin resistance cassette with UL34 flanking sequence was constructed in several steps. First a GmR cassette containing the GmR promoter, protein coding sequence and terminator flanked at the 5’ end with a SceI homing nuclease site and at the 3’ end by ORF24 and UL34 sequences were amplified from pEGFP-SceI-GmR using the primers Sce-GmR FW/ and ORF24 GmR RV1 (Table 3). Second, a PCR product containing the entire FLAG-tagged ORF24 flanked by UL34 flanking sequences at one end and GmR cassette homology at the other was amplified from VZV-infected cell DNA template using the primers FLAG-ORF24 FW and ORF24 Overlap RV. The two resulting PCR products overlap at one end of the GmR cassette sequence. The complete insertion cassette was then assembled in a PCR reaction with the overlapping PCR products as template and using the primers FLAG-ORF24 FW and ORF24 GmR RV2. The resulting PCR product was recombined into the HSV-1 (F) BAC, and Gm-resistant recombinants were picked, and genomes were tested for insertion of the Gm cassette by diagnostic PCR using the flanking primers ORF24 Diag FW and ORF24 Diag RV. Scarless excision of the Gm cassette, leaving a full FLAG-ORF 24 replacing most of the UL34 coding sequences was carried out, and Gm-sensitive clones were tested for correct structure by diagnostic PCR using the ORF24 Diag FW and ORF24 Diag RV primers. Proper insertion and excision were confirmed by Sanger sequencing of the UL34/ORF24 locus using the entire BAC DNA as template.

**Table 3.**
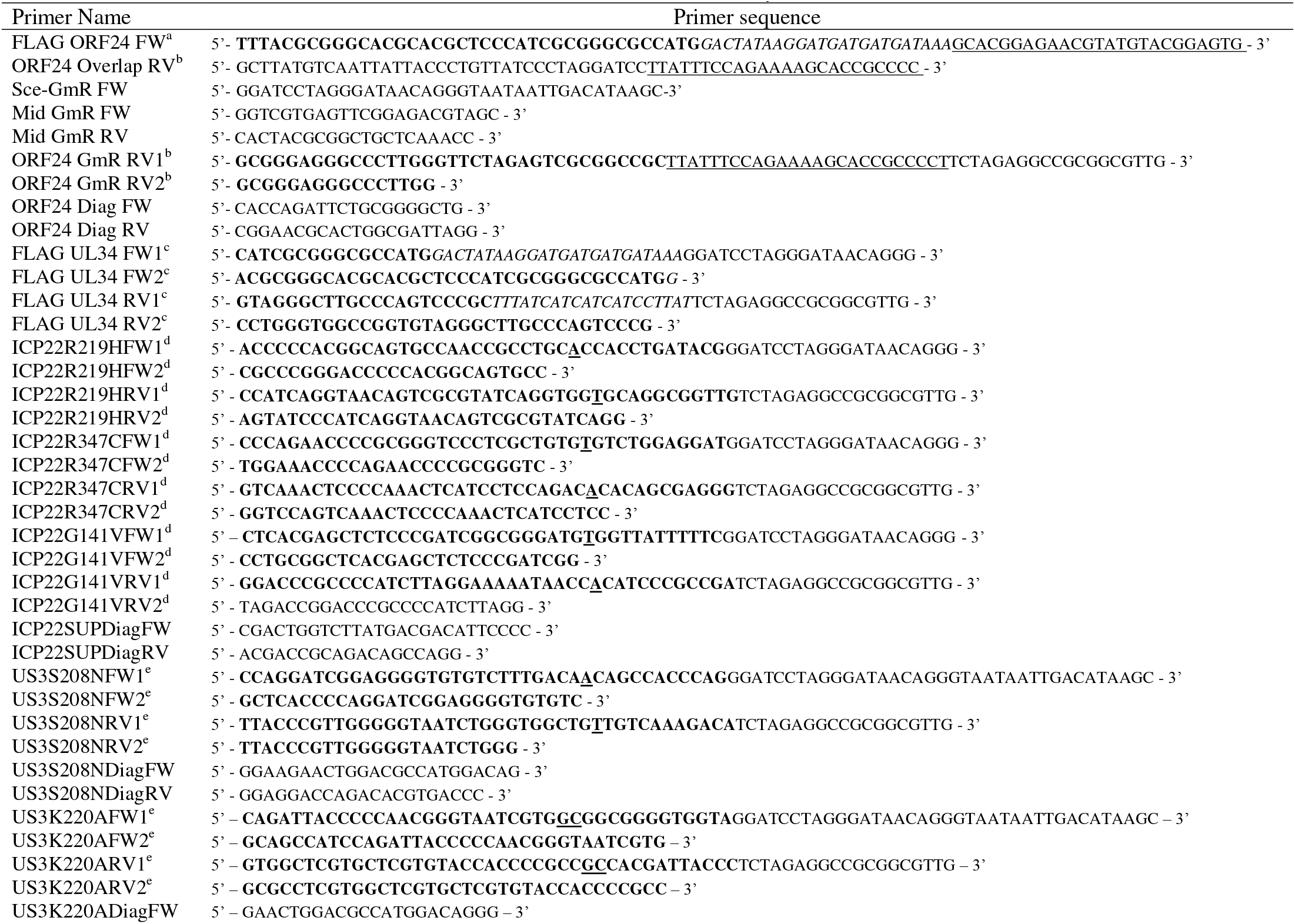
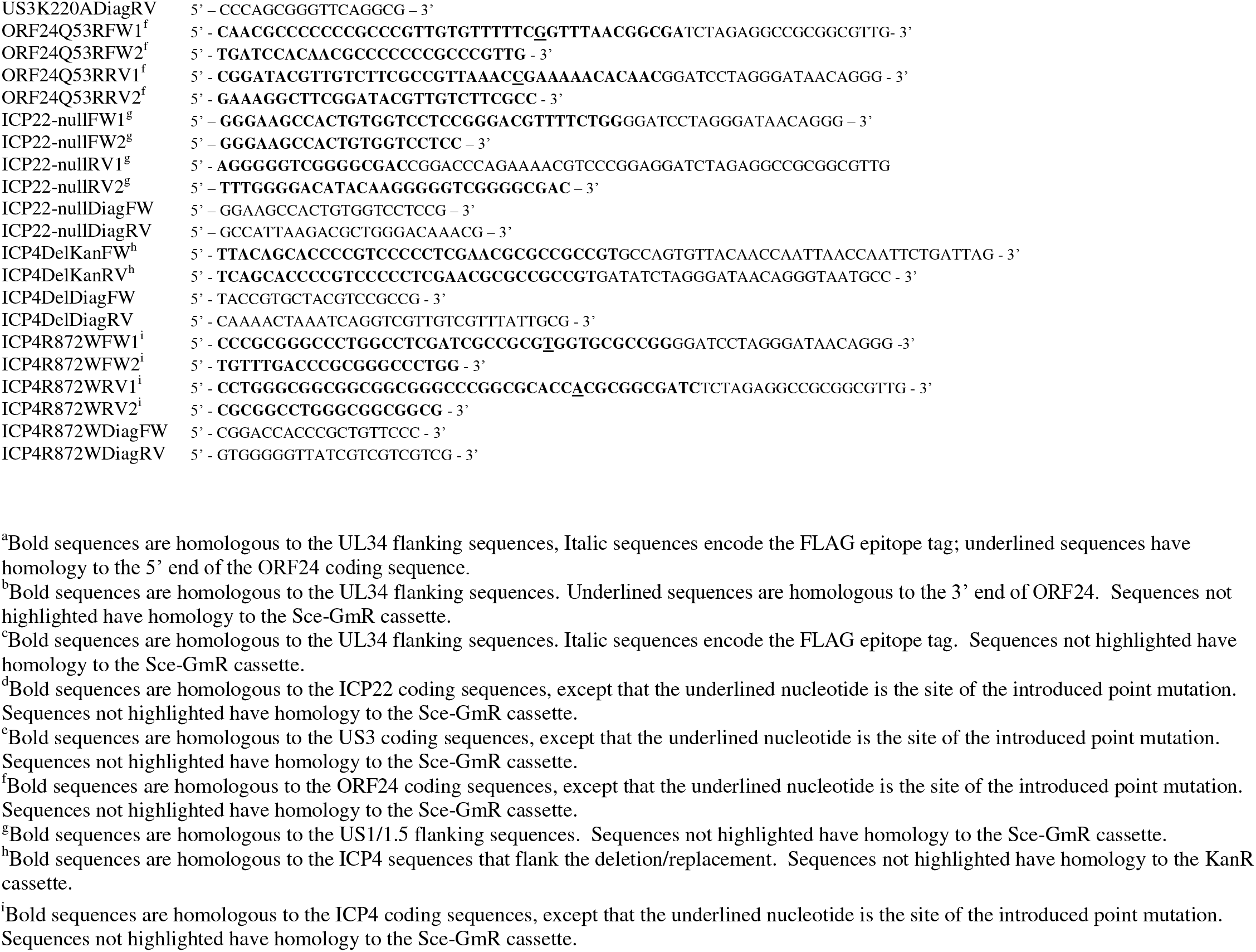
Primers used for construction and analysis of mutant BACs

For construction of an HSV-1 recombinant expressing FLAG-tagged pUL34 (referred to as FLAG-HSV-34), a cassette containing the FLAG epitope tag coding sequence, a GmR resistance gene and UL34 flanking sequences was constructed in two steps. First, PCR products containing the 5’ and 3’ halves of the Sce-GmR cassette were amplified from the pEGFP-Sce-GmR using the primers FLAG UL34 FW1 and Mid GmR RV, and Mid GmR FW and FLAG UL34 RV1, respectively (Table 3). The two resulting PCR products overlap in the Gm coding sequence. The complete mutagenic cassette was then assembled in a PCR reaction using the overlapping partial genes and the primers FLAG UL34 FW2 and FLAG UL34 RV2 (Table 1). The resulting PCR product was recombined into HSV-1(F) BAC and Gm-resistant recombinants were picked and genomes were tested for insertion of the Gm cassette by diagnostic PCR using flanking primers. Correct insertion of the Gm cassette was confirmed by direct sequencing of the BAC DNA. Scarless excision of the Gm cassette, leaving an intact FLAG-UL34 gene was carried out as described previously and Gm-sensitive clones were tested for correct structure at the UL34 locus by diagnostic PCR and correct structure in-frame insertion of the tag was confirmed by Sanger sequencing of the UL34 locus using the BAC DNA as template. Viruses were rescued from the chimeric BAC by transfection into UL34-expressing complementing cells. Correct incorporation of mutations into the final viruses was confirmed by PCR amplification and sequencing of the UL34 locus from the recombinant virus stock.

HSV recombinants encoding individual ORF24_Q53R_, ICP22_R219H_, ICP22_R347C_, ICP22_G141V_, and US3_S208N_ suppressor point mutations and the US3 catalytically inactivating point mutation US3_K220A_ were made on the FLAG-VZV-24 backgrounds by previously described methods for introducing point mutations (25) and using the primer sets in Table 3. Correct incorporation of mutations into the final viruses was confirmed by PCR amplification and sequencing of the mutant locus from the recombinant virus stock and by whole genome sequencing of the resulting virus genomes.

Scarless deletion of the US1/1.5 gene was performed as previously described using the primer sets in Table 3.

An HSV recombinant containing the ICP4 suppressor point mutation was made in two steps on the FLAG-VZV-24 background. First, one copy of the ICP4 gene in the FLAG-VZV-24 BAC was deleted and replaced by a kanamycin resistance cassette that had been amplified using the primers ICP4DelKanFW and ICP4DelKanRV shown in Table 3. Proper deletion and replacement of one of the ICP4 copies was demonstrated in two diagnostic PCR reactions, one using ICP4DelDiagFW and ICP4DelDiagRV to demonstrate replacement of one ICP4 coding sequence with the kanamycin resistance cassette, and the other using ICP4R872WDiagFW and ICP4R872WDiagRV to demonstrate the continued presence of ICP4 coding sequence at the second ICP4 coding sequence. On this background, the point mutation was introduced into the remaining ICP4 coding sequence using the same strategy for introduction of point mutations described above and using the ICP4R872WFW1, ICP4R872WFW2, ICP4R872WRV1, and ICP4R872WRV2 primers shown in Table 3. Presence of the ICP4 point mutation was confirmed in both the BAC and the subsequently rescued virus by Sanger sequencing of the PCR product generated using the ICP4R872WDiagFW and ICP4R872WDiagRV primers and by whole genome sequencing of capsid DNA isolated from cells infected with the recombinant virus.

### Immunoblotting

Nitrocellulose sheets bearing proteins of interest were blocked in 5% non-fat milk plus 0.2% Tween 20 for at least 2 h. The membranes were probed either with a mouse anti-FLAG M2 monoclonal antibody (Sigma/Aldrich) 1:1000, anti-HA mouse monoclonal antibody (BioLegend), mouse monoclonal anti-ICP27 (Virusys) 1:1000, or mouse anti-VP5 (Biodesign. International) 1:500. followed by reaction with alkaline phosphatase-conjugated secondary antibody.

### Plaque size assays for virus spread

Six-well tissue culture plates were seeded with 1.5 × 10^6^ Vero or UL34-complementing cells the day before infection. Cells were infected at a low MOI for 1.5 h, after which the virus inoculum was replaced with a 1:250 dilution of pooled human immunoglobulin (GammaSTAN) in infection medium. Cells were fixed and stained after 2 days of infection. A mouse monoclonal anti-HSV 45-kDa protein (scaffolding protein) antibody (Serotec) at 1:2000 dilution and Alexa Fluor 568–goat anti-mouse IgG (Invitrogen) at 1:1000 dilution were used as primary and secondary antibodies, respectively. Plaque images were outlined in ImageJ using the freehand tool and the enclosed area in pixels was measured.

### Single step growth assays

Measurement of replication of recombinant viruses on Vero, and UL34-complementing CX cells after infection at high multiplicity was performed as previously described (25).

### Indirect immunofluorescence

Immunofluorescence for co-localization of epitope tagged proteins was performed as previously described using either 1:500 anti-FLAG rabbit antiserum, 1:1000 M2 anti-FLAG mouse monoclonal antibody (SIGMA/Aldrich), and 1:500 anti-HA mouse monoclonal antibody (BioLegend) (58, 59). Alexa-tagged secondary antibodies, Hoechst 333342, and Alexa488-tagged phalloidin were obtained from Invitrogen and were used at 1:1000 dilution.

### Plasmid constructions

pcDNA plasmids carrying wt UL34 (pRR1238) or UL31-FLAG (pRR1334) were described previously (36). A pcDNA3 plasmid carrying N-terminally HA-tagged UL34 (pRR1385) was constructed by the amplification of the UL34 coding sequence from the HSV-1(F) BAC using primers 5′ – GATCAAGCTTCCATGTACCCATACGATGTTCCAGATTACGCTGCGGGACTGGGCAAGCCC—3′ and 5′ – CTAGTCTAGATTATAGGCGCGCGCCAGC—3’, digestion of the resulting PCR product with HindIII and AflII, and ligation into HindIII-AflII-cut pRR1238. Construction of a pcDNA3 plasmid encoding N-terminally HA-tagged ORF24 (pRR1408) was by Gibson assembly using a vector product derived from pcDNA3 and an ORF24 insert amplified from the ORF24 replacement BAC. The ORF24-HA insert was amplified using the primers 5’ – ATGTATCCATATGACGTCCCAGACTCTGCCGCACGGAGAACGTATGTACGGAGTG – 3’ and 5’ – ACTGGCGGCCGTTACTAGTGGATCCTTATTTCCAGAAAAGCACCGCCCC – 3’. The vector sequences were amplified using the primers 5’ – GGCAGAGTCTGGGACGTCATATGGATACATGAGCTCGGTACCAAGCTTGGGTC – 3’ and 5’ – GGATCCACTAGTAACGGCCGCC – 3’. The amplified PCR products were assembled using New England Biolabs 2X Gibson master mixing according to the manufacturer’s instructions.

### Co-immunoprecipitation assays

293T cells in 150 mm dishes were transfected with plasmids pRR1385 (HA-UL34), pRR1408 (HA-ORF24) pRR1334 (UL31-FLAG), or pcDNA3 using polyethyleneimine (PEI). A total of 12 μg of plasmid was diluted into 1 μl DMEM with no additives (DMEM/NA) and then mixed with 1 ml of DMEM/NA containing 108 μg PEI. After incubation at room temperature for 15 minutes to allow complexes to form, the mixture was added to cultures of 50% confluent cells that had been seeded 24 hours previously. After 48 hours, transfected cells were washed once in phosphate buffered saline (PBS) and then scraped into 5 ml fresh PBS and pelleted at 800xg for 10 minutes. Cells were lysed by resuspension of the cell pellet in RIPA buffer (50mMM Tris pH 7.5, 150 mM NaCl, 1 mM EDTA, 1% Triton X-100, 5 mM Na Vanadate, 5 mM NaF) followed by sonication. After removal of a 60 μl input control sample. The remainder was immunoprecipitated using 4 μl anti-HA magnetic resin resin (Sigma) according to the manufacturer’s instructions, using an overnight binding step and elution with hot SDS-PAGE sample buffer. Input and immunoprecipitation samples were separated in 10% SDS-PAGE gels and blotted onto nitrocellulose membranes. The scale of transfection and amount of magnetic resin used were optimized so that the resin was limiting for purification of HA-tagged protein. Using this approach, equivalent amounts of HA-pUL34 and HA-ORF24 were immunoprecipitated even though expression of HA-ORF24 was invariably lower than HA-pUL34.

### Selection and amplification of phenotypic suppressor viruses

Six parallel 5 cm^2^ cultures of Vero cells were each infected with 1000 PFU of VZV ORF24 #1 virus and allowed to grow for 48 hours, by which time only minute clusters of infected cells were observed. A virus stock was then prepared from each culture and half of it was used to infect a fresh 25 cm^2^ culture of Vero cells, and infection was allowed to proceed for an additional two days by which time extensive CPE was evident in all of the cultures. A virus stock was prepared from each of the cultures and a single virus was plaque purified on Vero cells through two rounds from each of the six selection stocks. The resulting plaque purified viruses were amplified to high titer stocks on HSV UL34-expressing complementing cells.

### Preparation of viral DNA for genome sequencing

Vero or UL34 complementing cells in 100 mm tissue culture plates were infected with 5 PFU/cell of virus in 3 ml infection medium (DMEM high glucose supplemented with 1% heat inactivated calf serum) for 90 minutes and then the inoculum was replaced with 10 ml fresh infection medium and infection was allowed to proceed for 20 hours. The infection medium was then removed and cells were washed with 5 ml phosphate buffered saline (PBS) and then scraped into 5 ml of fresh PBS and transferred to a 15 ml conical centrifuge tube. Cells were pelleted at 300x*g* for 5 minutes and the cell pellet was then resuspended in 1 ml PBS containing 0.5% nonidet P-40 (NP-40) and frozen at −80° C. The cell lysate was then thawed and sonicated twice for 20 seconds. Cell debris was pelleted in the microcentrifuge at 10,000x*g* for 3 minutes. The supernatant liquid was transferred to a fresh tube and 10 μl of 1 M MgCl2, 4 μl of 250X protease inhibitor cocktail (SIGMA), 100 U DNase I (Worthington), and 50 μg RNase A (SIGMA) were added and the reaction was incubated at 30° C for 30 minutes to degrade non-encapsidated nucleic acids. Debris that precipitated during the digestion was removed by centrifugation again in the microcentrifuge at 10,000x*g* for 3 minutes. The supernatant liquid was transferred to a fresh microcentrifuge tube and capsids were pelleted by centrifugation at 21,000x*g* for 1 h. The supernatant liquid was discarded and the capsid pellet was resuspended by repeated pipetting in 250 μl PBS. EDTA and SDS were added to final concentrations of 5 mM and 1 %, respectively and the mixture was incubated at 37° C for 30 minutes to release DNA from the capsids. DNA was purified by phenol/chloroform extraction and ethanol precipitation.

### Genome sequence analysis

Illumina library preparation and genome sequencing on a NextSeq 2000 platform was performed by the Microbial Genome Sequencing Center Pittsburgh, PA. A consensus sequence for the ORF24 #1 BAC was derived by alignment of unpaired-end Illumina reads derived from the BAC DNA to an *in silico* constructed reference sequence built using information from the published sequence of HSV-1(F), the published sequence of the BAC vector sequences used for construction of the HSV-1(F) parental BAC, and the sequence of ORF24 derived from Sanger sequencing of the PCR product used for mutagenesis (32, 33). Alignment was performed using Bowtie implemented on the Unipro UGENE platform using the default alignment settings (60, 61). Consensus sequences of the ORF24 #1 parental virus genome and of the suppressor variants were derived by alignment of Illumina reads with the experimentally determined ORF24#1 BAC sequence. The consensus sequence of the parental virus genome was identical to that of the BAC from which it was derived.

### Transmission electron microscopy

Confluent monolayers of Vero cells were infected at an MOI of 5 in infection medium for 20 hours and then fixed by incubation in 2.5% glutaraldehyde in 0.1 M cacodylate buffer (pH 7.4) for 2 h. Cells were postfixed in 1% osmium tetroxide, washed in cacodylate buffer, embedded in Spurr’s resin, and cut into 95-nm sections. Sections were mounted on grids, stained with uranyl acetate and lead citrate, and examined with a JEOL 1250 transmission electron microscope. Ten cells were used for the quantitation of viral capsids within cellular compartments. DNA- and non-DNA-containing capsids were counted in the nucleus, the perinuclear space (between the INM and the ONM), and the extranuclear compartment (within the cytoplasm or the extracellular space).

### Nuclear contour assays

Cross-sectional TEM images of nuclei of infected Vero cells obtained as described above were used to measure the nuclear contour ratio. The nuclear cross-sectional area and perimeter from 1000X magnified images were measured using ImageJ and the nuclear contour ratio was calculated using the formula (4π × area)/perimeter^2^, as described by Smeulders and Dorst (62). Twenty images of each sample were measured.

## Acknowledgments

The authors are grateful to Yasushi Kawaguchi and Greg Smith for providing reagents for recombinant virus construction, and to Shaowen White for critical reading of the manuscript. This work was funded by NIH grants R21AI133155, R21AI148831, and R21AI153683.

